# Cell type identity determines transcriptomic immune responses in *Arabidopsis thaliana* roots

**DOI:** 10.1101/302448

**Authors:** Charlotte Rich, Marco U. Reitz, Ruth Eichmann, Sophie Hermann, Dafyd J. Jenkins, Karl-Heinz Kogel, Eddi Esteban, Sascha Ott, Patrick Schäfer

## Abstract

Root pathogens are a major threat in global crop production and protection strategies are required to sustainably enhance the efficiency of root immunity. Our understanding of root immunity is still limited relative to our knowledge of immune responses in leaves. In an effort to reveal the organisation of immunity in roots, we undertook a cell type-specific transcriptome analysis to identify gene networks activated in epidermis, cortex and pericycle cells of *Arabidopsis* roots upon treatment with two immunity elicitors, the bacterial microbe-associated molecular pattern flagellin, and the endogenous damage-associated molecular pattern Pep1. Our analyses revealed that both elicitors induced immunity gene networks in a cell type-specific manner. Interestingly, both elicitors did not alter cell identity-determining gene networks. Using sophisticated paired motif promoter analyses, we identified key transcription factor pairs involved in the regulation of cell type-specific immunity networks. In addition, our data show that cell identity networks integrate with cell immunity networks to activate cell type-specific immune response according to the functional capabilities of each cell type.

**Material Distribution Footnote:** The author responsible for distribution of materials integral to the findings presented in this article in accordance with the policy described in the Instructions for Authors (www.plantcell.org) is: Patrick Schäfer.

## Introduction

Plant roots are essential for plant health and development. In addition to anchoring plants in the ground, roots capture nutrients and water, and provide protection from soil-based microbes. Conducting these different tasks is especially challenging under changing environments and acute stress conditions. Roots have evolved complex tissues comprising a diversity of cell types with different functions that govern overall root functionality and provide the necessary plasticity to cope with environmental stress. Organised in concentric layers, *Arabidopsis* roots consist of an outermost epidermis, followed by cortex, endodermis, pericycle and the root vascular tissue with xylem and phloem cells (Dolan et al., 1993; Brady et al., 2007). This organisation is implemented by the stem cell niche in the very root tip where cell fate is determined and cell types maintain their given identity throughout their lifetime (van den Berg et al., 1995; Sabatini et al., 2003; Wendrich et al., 2017). Overall, the concerted interplay of different cell type functions ascertain root functionality and plasticity (Brady et al., 2007; Birnbaum, 2003; Walker et al., 2017). Understanding this inter-dependency of cell types and its accurate spatio-temporal regulation is of particular relevance for our efforts to generate crops with improved resilience under stressful environments.

Studies of cell type-specific transcriptomics based on fluorescence-activated cell sorting have significantly advanced our knowledge of processes regulating root development (Birnbaum, 2003; Birnbaum et al., 2005; Bargmann et al., 2013; Walker et al., 2017) and furthered our understanding of root adaptation to abiotic stress and nutrient depletion (Dinneny et al., 2008; Gifford et al., 2008; Geng et al., 2013; Gifford et al., 2013). Evidence provided in these studies highlighted the functional individuality of cell types and the significance of a coordinated regulation of cell type-specific gene networks to master root development and secure overall root functionality (e.g. growth) under stress.

While root cell type-specific transcriptomics have addressed key questions in root development and abiotic stress adaptation, the function of root cell types in regulating root immunity remains elusive. Root diseases represent a major threat in crop production and enhancing root resistance against pathogens by improving processes regulating pattern-triggered immunity (PTI) is of outstanding importance. The concept of PTI emerged from studies in leaves. Plasma membrane-localised pattern recognition receptors (PRRs) recognise microbe-associated molecular patterns (MAMPs) as non-self molecules from microbes to induce PTI (Jones and Dangl, 2006; Boller and Felix, 2009). Leaves, for instance, elicited with the MAMP flg22 (the active epitope of bacterial flagellin, recognised by the PRR FLS2) activate PTI responses including the rapid production of reactive oxygen species (ROS burst), MITOGEN-ACTIVATED PROTEIN KINASE (MAPK) phosphorylation and induction of immunity genes to stop pathogen infection (Felix et al., 1999; Gómez-Gómez et al., 1999; Asai et al., 2002; Zipfel et al., 2004). Similarly, root cells recognise MAMPs such as flg22 by FLAGELLIN SENSITIVE 2 (FLS2) to trigger effective PTI (Millet et al., 2010; Jacobs et al., 2011; Beck et al., 2014; Wyrsch et al., 2015; Poncini et al., 2017). In addition to MAMPs, plants produce damage-associated molecular patterns (DAMPs) upon pathogen recognition as part of PTI. Among the DAMPs produced in *Arabidopsis* are Pep1-Pep8 peptides encoded by respective *PROPEP* genes (Flury et al., 2013). Pep1, encoded by *PROPEP1*, is one of the best studied Pep and is recognised by the PRRs PEP RECEPTOR 1 and 2 (PEPR1 and PEPR2) at the plasma membrane, inducing similar PTI responses as reported for flg22 (Huffaker et al., 2006; Yamaguchi et al., 2006; Krol et al., 2010; Yamaguchi et al., 2010). Consistently, PEPR1/2 and FLS2 pathways share signalling components such as MAPKs (Schulze et al., 2010; Liu et al., 2013; Yamada et al., 2016). Recent studies, however, indicate certain differences between MAMP (e.g. flg22) and Pep-induced PTI that might be at least partially explained by additional activities of PEPRs. Qi et al. (2010) identified a unique guanylyl cyclase activity for PEPR1 to activate CYCLIC NUCLEOTIDE-GATED CHANNEL 2 (CNGC2)-mediated apoplastic Ca^2+^ influx as part of Pep-induced PTI signalling. PEPR1/2-triggered immune signalling was further shown to maintain PTI in plants lacking MAMP perception and signalling components (Tintor et al., 2013; Yamada et al., 2016). Accordingly, comparative analyses of flg22 and Pep1 activity in roots, while demonstrating typical PTI responses, Pep1 was a more potent elicitor of immunity (Poncini et al., 2017). While all these studies highlight apparent differences and interdependencies between flg22 and Pep1-induced PTI in *Arabidopsis* the gene network defining DAMP and MAMP-mediated PTI in roots is currently unknown.

Motivated by recent findings suggesting distinct competences of different root cell types in launching PTI (Wyrsch et al., 2015), we studied the contribution of different root cell types to PTI activation. Here we describe the transcriptional networks of three *Arabidopsis* root cell types; epidermis, cortex and pericycle, following treatment by flg22 and Pep1. Our study demonstrates that different immunity gene networks are activated in the three cell types and, hence, these cell types contribute differently to overall root PTI. To explain different transcriptomic responses, we particularly focussed on transcription factors (TFs) and combinatorial TF motif analyses. Activated by signalling cascades TFs combinations provide regulatory specificity to orchestrate transcriptomic responses. By developing a statistical test for enrichment of paired TF motifs that accounts for multiplicity of binding sites, overlapping sites, and weak binding sites, we were able to explain cell type-specific differences in immune responses by specific TF motif combinations. We further identified differences between Pep1 and flg22-activated gene networks and the associated combinations of TFs on the cell type level. Moreover, we explored the connection of cell identity and immunity-determining gene networks and discuss the significance of their self-contained co-existence in specifying cell type functionality and, hence, securing root integrity under conditions of environmental stress.

## Results

### flg22 and Pep1 activate root immunity through partially different signalling pathways

Treatment of *Arabidopsis* roots by immunity elicitors flg22 or Pep1 induces hallmark PTI responses, including a ROS burst (Figure 1A), activated MAPK signalling cascades as evidenced by MAPK phosphorylation (Figure 1B) and induction of immunity genes in roots (Figure 1C and D). Activated PTI eventually leads to plant growth inhibition (Figure 1E and F). flg22 and Pep1 have been proposed to act through overlapping pathways (Krol et al., 2010; Yamaguchi and Huffaker, 2011; Tintor et al., 2013). We previously demonstrated that the beneficial root endophyte *Serendipita indica* (formerly *Piriformospora indica*) suppressed PTI to facilitate root colonisation (Jacobs et al., 2011). When applying *S. indica* to *Arabidopsis* roots, the fungus inhibits MAPK phosphorylation, PTI marker gene induction and growth inhibition that occurs following flg22 treatment (Fig. 1B, C, E, Suplemental Figure 1A). Interestingly, *S. indica* did not suppress the Pep1-induced ROS burst, MAPK phosphorylation nor PTI marker gene expression (Figure 1A and F, Supplemental Figure 1A, B, D) and could not rescue Pep1-induced growth inhibition (Figure 1 F). We previously showed that flg22 treatment of roots inhibited *S. indica* colonisation (Jacobs et al., 2011). Consistently, genetic evidence supporting effective Pep1-induced immunity was illustrated by enhanced *S. indica* colonisation of roots in the Pep1 receptor mutant *pepr1 pepr2* (Supplemental Figure 1C). These data therefore suggest that flg22 and Pep1 recruit, at least partially, different signalling pathways to activate PTI in roots. Considering the diverse functions of root cell types in root development (Birnbaum et al., 2003; Brady et al., 2007; Gifford et al., 2013) and cell type-specific gene expression networks upon abiotic stress (Dinneny et al., 2008; Geng et al., 2013), we next explored to what extent root cell types differed in PTI transcriptional by performing RNA-seq on three root cell type populations following treatment with flg22 or Pep1.

**Figure 1.**
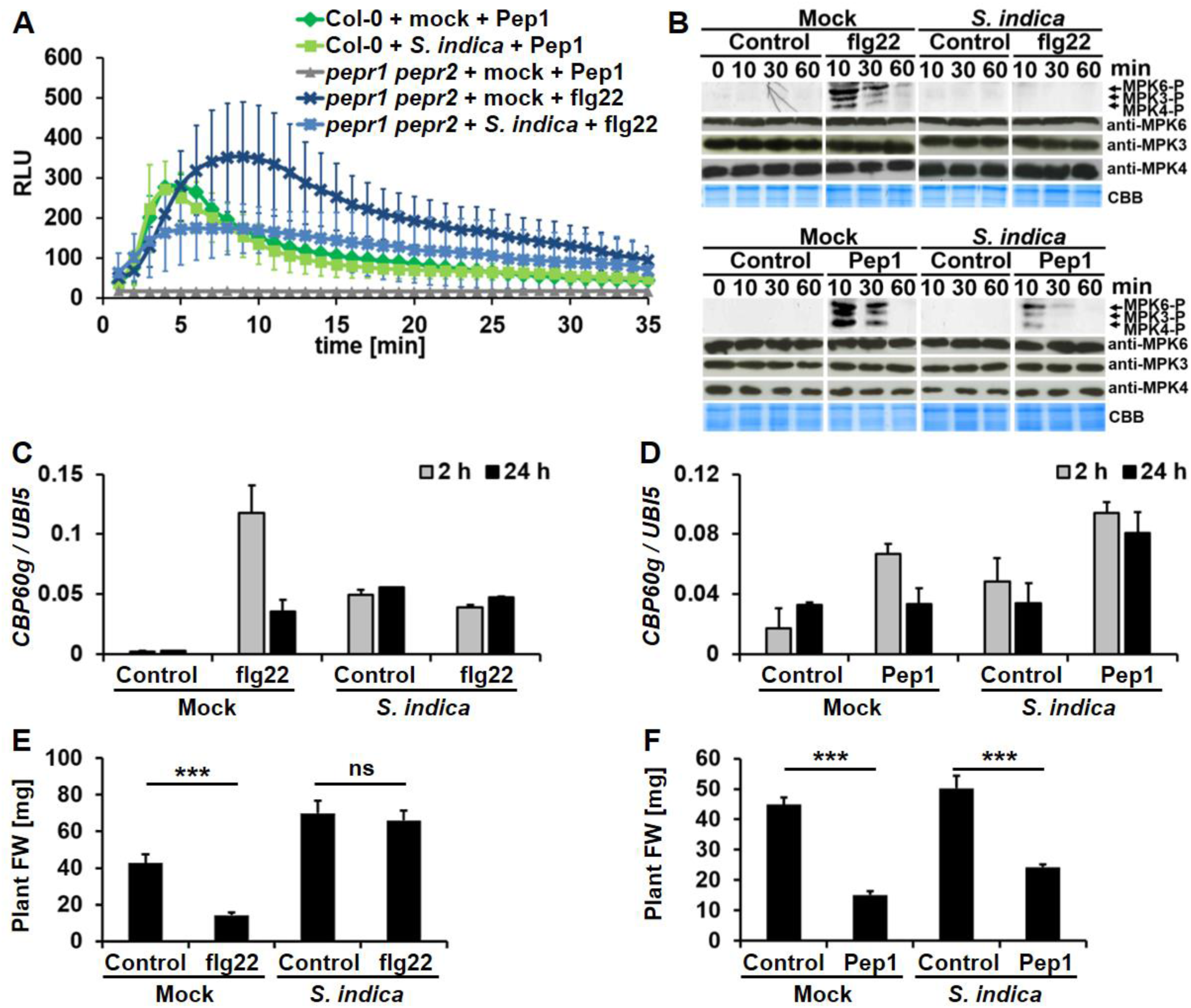
The ability of *Serendipita indica* to suppress flg22 but not Pep1-triggered immune responses in roots. **(A)** Pep1-induced oxidative burst in roots is dependent on PEPR1 and PEPR2 and, in contrast to flg22-induced oxidative burst, cannot be suppressed by *S. indica*. RLU, relative light units. **(B)** *S. indica* inhibits flg22 and only partially Pep1-induced MPK phosphorylation (MPKx-P) in roots. Note that the abundance of MPKs is not affected by any treatment. CBB, coomassie brilliant blue. **(C, D)** *S. indica* inhibits flg22 but not Pep1 induction of PTI marker gene *CBP60g*. h, hours after flg22 or Pep1 treatment. **(E, F)** *S. indica* abolishes flg22 but not Pep1-induced seedling growth inhibition. FW, fresh weight; ***, p < 0.001; ns, not significant.

### Root cell types differ in their immunity gene networks

Separated by the lignified and suberized endodermis, roots are divided into outer cell layers (epidermis, cortex), which are in contact with the soil environment, and inner cell layers forming the water and nutrient transporting vasculature. Representing the interface with the soil environment, the epidermis and cortex are expected to be crucial for root protection against pathogen invasion. As pericycle cells, the outermost cell layer of the inner root tissue, guard the vasculature and were recently shown to be highly responsive to immune elicitors (Wyrsch et al., 2015). Therefore, pericycle, epidermis and cortex cells were selected for our transcriptional analyses. To determine if immune signalling networks differ between cell types, we treated roots of *Arabidopsis* lines specifically expressing GFP in epidermis (*pGL2:GFP*), cortex (*pCORTEX:GFP*) or pericycle (*E3754*) (Brady et al., 2007; Gifford et al., 2008) with either flg22 or Pep1. After protoplasting, GFP-expressing cells were extracted using fluorescent-activated cell sorting (FACS; Birnbaum et al. 2003; Brady et al. 2007) (Figure 2A). As reported for cell type transcriptome analyses of abiotic stress networks (Dinneny et al., 2008; Geng et al., 2013), we treated whole roots rather than protoplasts to ensure the tissue context and intercellular communication of cell types were preserved. Effective PTI depends on fast activation of immunity pathways with the most essential immune responses (e.g. ROS burst, MAPK phosphorylation, transcriptional reprogramming) activated within the first two hours after pathogen recognition (Zipfel et al., 2004, Fig. 1). We therefore focused on the response at 2 h post elicitor treatment to capture the initial primary differences between flg22 and Pep1-triggered immunity pathways, prior to secondary activation of cross-talking networks and later PTI responses such as development and growth-related gene networks as a result of PTI-induced growth inhibition (observed at 4 d post elictor treatment. In addition, previous studies have demonstrated PEPR1/2 and FLS2 protein abundance in these cell types and the fast movement of flg22 and Pep1 reaching pericycle cells within a few minutes (Wyrsch et al., 2015; Ortiz-Morea et al., 2016).

**Figure 2.**
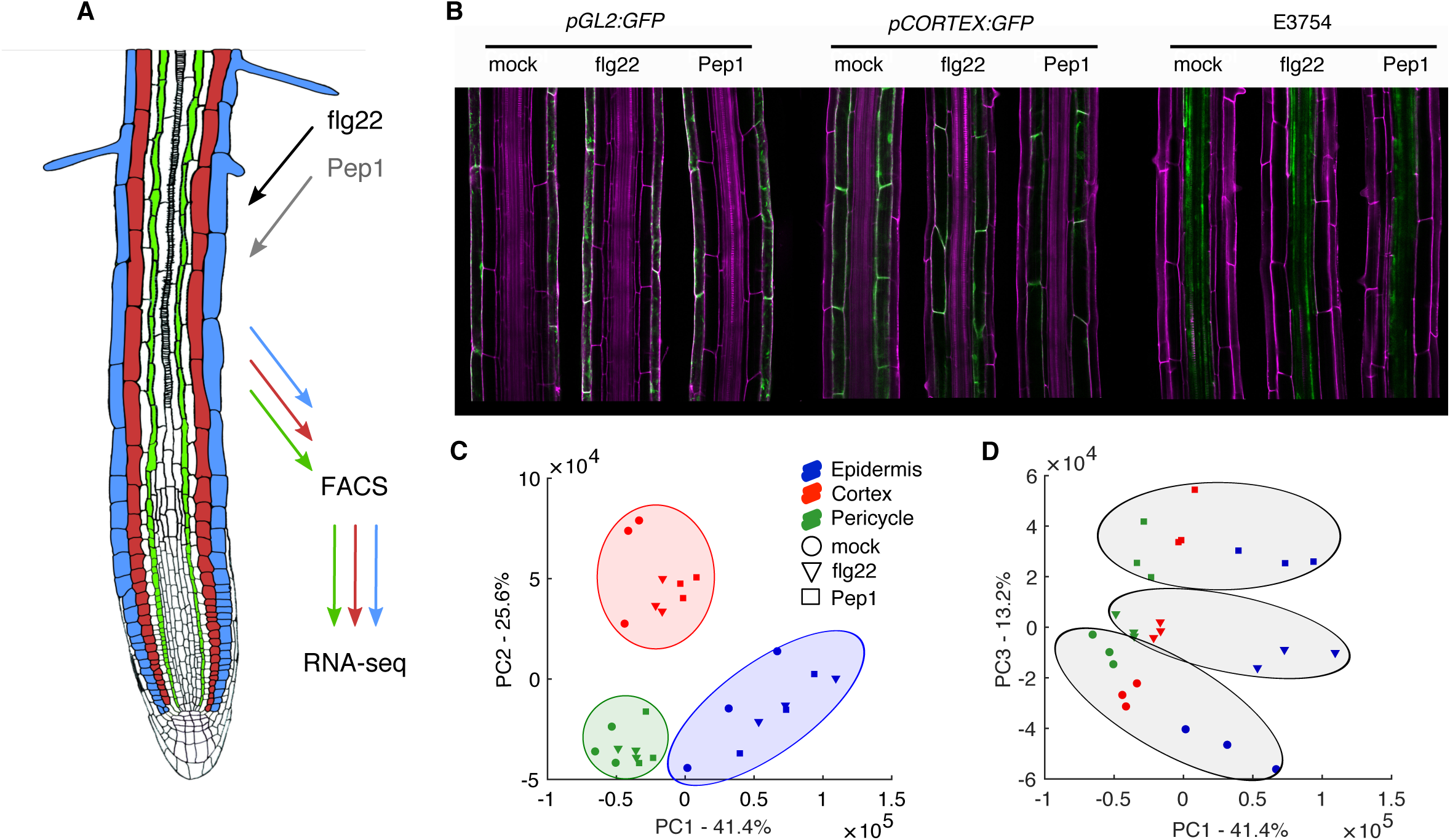
Measurement of transcriptomic responses to immune elicitors in three cell types. **(A)** Schematic showing experimental design. **(B)** Confocal images of cell-type specific markers *pGL2:GFP* (epidermis), *pCORTEX:GFP* (cortex) and E3754 (pericycle). **(C) and (D)** Principal component analysis of all replicates for each cell type and treatment comparing PC1 and 2 in (C), and PC1 and 3 in (D).

We first confirmed that cell type marker gene expression was unchanged in immunity activated plants by using confocal laser-scanning microscopy (Figure 2B) ensuring uniformity of our isolated cell populations. 100-bp paired RNA-seq reads were generated from these isolated cells (Figure 2A) and approximately 315 million reads (an average of 11.6 million read pairs per library) were uniquely mapped to gene features in the *Arabidopsis* genome (TAIR 10 assembly, see Materials and Methods, Supplemental Table 1). Principal Component Analysis (PCA) of the RNA-seq samples revealed the majority (∼80%) of the variation within the dataset was contained within the first three principal components (Supplemental Figure 2A). Cell identity was the principal source of variation, as principal components (PCs) 1 (41.4%) and 2 (25.6%) clearly clustered by cell type (Figure 2C and D). Consistent with this, cell type marker genes were highly expressed in the respective cell type populations (Supplemental Figure 2C-E). A second major source of variation (plotting PC 1 against PC3 (13.2%)) suggested that root cell types responded differently to flg22 and Pep1 (Figure 2D, Supplemental Figure 2A).

Next, we analysed our RNA-seq data to identify differentially expressed gene (DEG) sets in each cell type in response to flg22 and Pep1 elicitation. A total of 3,159 unique genes were differentially expressed in response to one or both elicitors in at least one cell type (Figure 3A). Consistent with a recent study (Poncini et al., 2017), Pep1 treatment elicited the most DEGs (2988), with approximately equally numbers of up- and down-regulated genes, whereas much fewer DEGS (817) were elicited by flg22 treatment. Collectively, these data support the hypothesis that Pep1 activates different transcriptional networks to flg22. Focussing on the DEGs in the specific cell types, 573 (epidermis), 434 (cortex) and 64 (pericycle) genes were differentially expressed by flg22 treatment whereas DEGs elicited by Pep1 were 1,679 (epidermis), 2,158 (cortex) and 360 (pericycle) genes (Figure 3A, Supplemental Table 2, Supplemental Data Set 1). We noted that the pericycle replicates were less consistent and noisier than the other two cell types, which may have contributed to the reduced number of DEGs observed (Supplemental Figure 4). Alternatively, a reduced number of DEGs may indicate that PTI is not consistently activated in root core tissue at this early time point (Supplemental Figure 4E-F, Supplemental Table 2). Interestingly, 74% (608 genes) of all flg22 and 66% (1,985 genes) of all Pep1-responsive genes were expressed in a single cell type (Figure 3B and C, Supplemental Data Set 2). By contrast, 26% of flg22 and 33% of Pep1-responsive genes were common to all cell types. Only 28 genes formed the “core” PTI response across all cell types following either flg22 or Pep1 treatments (Supplemental Figure 3, Supplemental Table 3). Within this core set we identified genes encoding two *GLUTATHIONE S-TRANSFERASES 1* and *11* (*GST1/11*), three O-methyltransferase family proteins, *INDOLE GLUCOSINOLATE METHYLTRANSFERASE 2, 3* and *4 (IGMT2/3/4*), various chitinases and peroxidases including *PEROXIDASE 4* and *71* (*PER4* and *PRX71)*. Taken together, our data show that both, flg22 and Pep1, activate different gene networks in each cell type.

**Figure 3.**
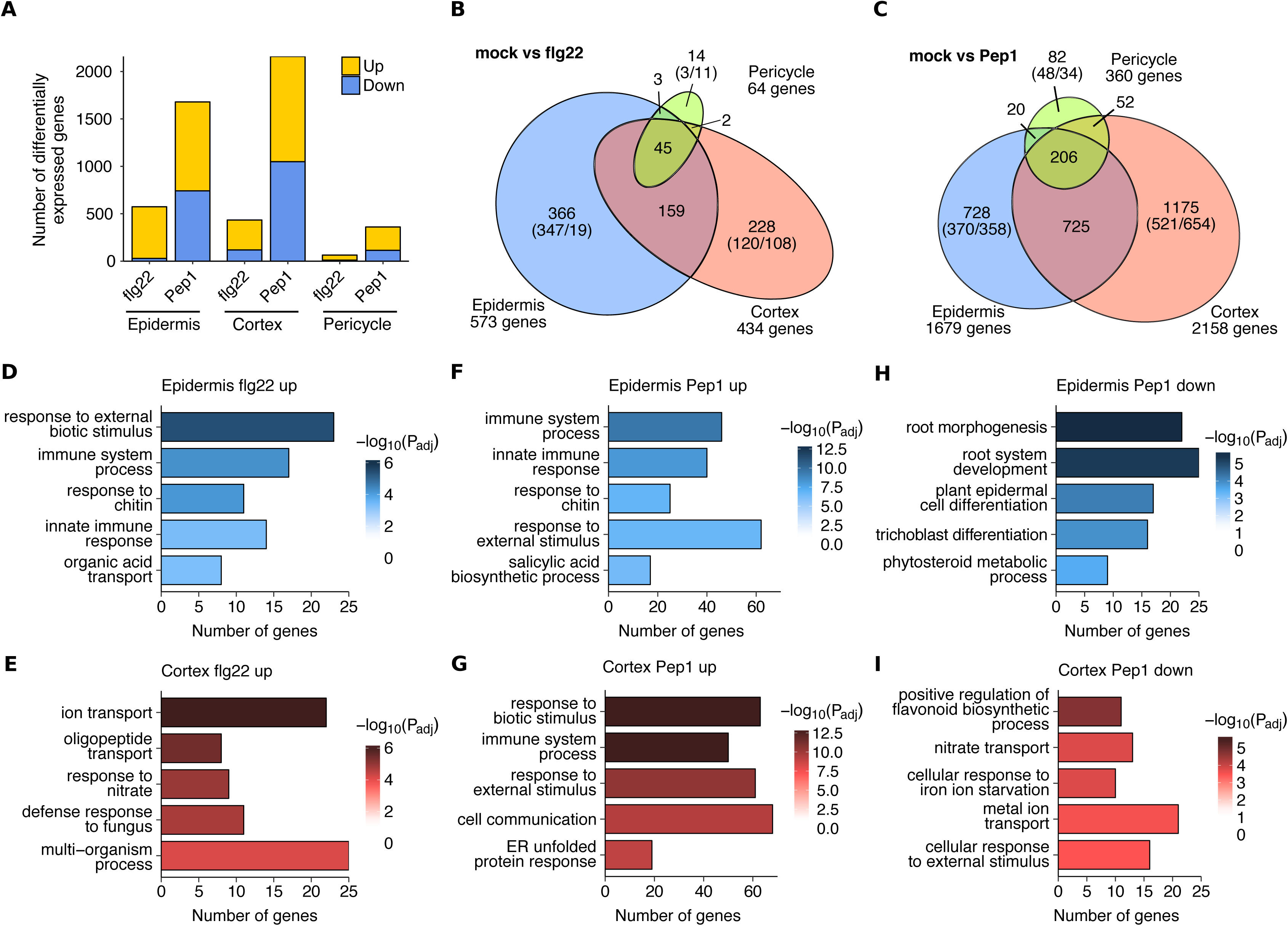
Root cell types differ in their immunity gene networks. **(A)** Numbers of differentially expressed genes in response to flg22 and Pep1 in different cell types. **(B)** Venn diagram showing cell type-specific networks responding to flg22. Numbers in brackets indicate up- and down-regulated genes. **(C)** Venn diagram showing cell type-specific networks responding to Pep1. Numbers in brackets indicate up- and down-regulated genes. **(D)** Top 5 most significantly enriched GO terms in genes up-regulated after flg22 treatment in the epidermis but not DE after flg22 treatment in the cortex or pericycle. **(E)** Top 5 most significantly enriched GO terms in genes up-regulated after flg22 treatment in the cortex but not DE after flg22 treatment in the epidermis or pericycle. **(F)**, **(G)**, **(H) and (I)** Top 5 most significantly enriched GO terms in exclusively up- or down-regulated genes after Pep1 treatment in the epidermis and cortex.

To explore if the observed cell type specificity of flg22 and Pep1-responsive gene networks reflects specific functions, we performed GO term enrichment analysis (Supplemental Table 4) on the epidermis and cortex datasets. To allow comparative GO term analyses we equalised gene set sizes for flg22 up-, Pep1 up- and Pep1 down-regulated genes (gene set sizes of 120 [cortex, flg22 up], 370 [epidermis, Pep1 up] and 358 [epidermis, Pep1 down] genes, respectively; see Figure 3B and C). Since relatively few DEGs were identified to be down-regulated by flg22 and overall in the pericycle, we investigated the function of these genes directly rather than by GO enrichment. Firstly, the up-regulated genes following both flg22 and Pep1 treatments in both the epidermis and the cortex were strongly enriched in many immunity-associated terms such as ‘immune system process’ and ‘regulation of defence response’ (Figure 3D-G). These terms are enriched in both the epidermis and the cortex despite the lack of overlap in the genes tested indicating that both cell types are immune-responsive via either temporally separated induction or represent immunity sub-networks.

In total, 21% (25/120) of epidermis-specific flg22-responsive genes were associated with immunity and defence terms annotated to genes such as *NDR1/HIN1-LIKE PROTEIN 10* (*NHL10*), *MITOGEN-ACTIVATED PROTEIN KINASE 5* (*MPK5*) and CHITINASE IV (*AtCHITIV*), and in the cortex, this proportion was 22.5% (27/120) of the cortex-specific DEGs including *CHITIN ELICITOR RECEPTOR KINASE 1* (*CERK1*), *WALL-ASSOCIATED RECEPTOR KINASE-LIKE 2* (*WAKL2*) and *WRKY8*. Similarly, Pep1 induced similar proportions of genes associated with immunity and defence; 18% in the epidermis (67/370) including *WRKY33, FLG22-INDUCED RECEPTOR-LIKE KINASE 1* (*FRK1*) and *PROPEP3*, and 19% in the cortex (70/370) including *BAK1-INTERACTING RECEPTOR-LIKE KINASE 1* (*BIR1*), *WRKY22* and *ARABIDOPSIS NAC DOMAIN-CONTAINING PROTEIN 019* (*ANAC019*). The DEG response to flg22 in the pericycle was limited to 14 largely uncharacterised genes with putative functions in protein modification and *CYP71A12* that functions in antimicrobial camalexin synthesis (Millet et al., 2010; Ludwig-Müller, 2015) (Supplemental Data Set 2).

In addition to the strong cell type-specific immune responses observed, GO enrichment analysis revealed functional specificity in both the epidermis and the cortex. ER stress associated terms such as ‘endoplasmic reticulum unfolded protein response’ (ER UPR) were more strongly enriched in the cortex (p<10^−4^ and p<10^−8^, cortex flg22 and Pep1 up-regulated DEGs, respectively) relative to their equivalent epidermis comparisons (p=1 and p<10^−3^ in the epidermis-specific flg22-and Pep1-responses, respectively) (Figure 3D and G). Genes associated with ER stress included *UDP-GLYCOSYLTRANSFERASE 85A1* (*UGT85A1*), *UDP-GALACTOSE TRANSPORTER 3* (*UTR3*) and the thiol-disulfide oxidoreductase *ENDOPLASMIC RETICULUM OXIDOREDUCTIN 1* (*ERO1*). Accordingly, leaf studies demonstrated the high relevance of antimicrobial synthesis and delivery for PTI effectivity (Kwon et al., 2008; Bednarek et al., 2009; Li et al., 2009; Nekrasov et al., 2009; Saijo et al., 2009). ‘Oligopeptide transport’ and ‘amide transport’ were specifically enriched in the cortex flg22- and Pep1-specific DEGs, associated to genes such as the sugar transporter *PROBABLE POLYOL TRANSPORTER* 6 (*PLT6*), *NITRATE TRANSPORTER 1.8* (*NRT1.8*) (flg22 response) and the sulphate transporter *SEEDLING LETHAL 1* (*SEL1*) (Pep1 response) (Figure 3E and G).

In the epidermis, up-regulated genes did not reveal a unique function as these genes were dominated by immunity-associated genes. However, the Pep1-repressed genes in the epidermis were more strongly enriched in development-associated GO terms such as ‘root morphogenesis’, ‘root system development’ and ‘plant epidermis cell differentiation’ (Figure 3H) than in other tested gene sets (p<10^−5^ in the epidermis Pep1 down-regulated versus p<0.03 in the cortex, ‘root morphogenesis’ term). This enrichment was associated with expansins such as EXPANSIN 14 (*EXP14*), auxin-associated gene *ANTHOCYANIN-IMPAIRED-RESPONSE 1* (*AIR1*) and development-associated gene KANADI 4 (*KAN4*). In contrast, the cortex down-regulated genes are uniquely enriched in terms associated with cell wall biogenesis such as ‘plant-type cell wall biogenesis’ (p<10^−4^) indicating that Pep1 could also be affecting development in the cortex via a functionally different set of genes (Figure 3I). Two expansins were also identified in the pericycle Pep1 down-regulated DEGs: *EXP15* and *EXPB3*. These combined results show that both flg22 and Pep1 are activating gene networks with distinct functions in all three cell types (Supplemental Data Set 2).

### The flg22 response is largely encompassed by the Pep1 response

In addition to cell type-specific patterns of GO terms, we also identified treatment-specific differences, particularly in the Pep1 response in both the epidermis and the cortex. For example, Pep1 activated genes associated with salicylic acid (SA)-associated GO terms including ‘SA biosynthetic process’ (Figure 3F) and ‘systemic acquired resistance’, ‘SA mediated signalling pathway’ much more strongly than flg22 (p<10^−6^ and p<0.03 in systemic acquired resistance in the epidermis Pep1- and flg22-response, respectively, Supplemental Data Set 3). SA associated genes include *WRKY33, CYSTEINE RICH RECEPTOR-LIKE PROTEIN KINASE 45* (*CRK45*) and *CYCLOPHILIN 38* (*CYP38*) in the epidermis, and *GLUTATHIONE S-TRANSFERASE TAU 7* (*GSTU7*) and *ANAC019* in the cortex. Pep1 additionally appears to impact brassinosteroid signalling with ‘brassinosteroid biosynthetic process’ enriched in both cortex and epidermis Pep1-repressed gene sets.

flg22 and Pep1 strongly overlapped in the enrichment of immunity and defence terms and we wanted to resolve the overlap and differences between flg22 and Pep1-induced DEGs in each cell type (Figure 4). Aggregating across cell types, ∼79% (646 of 817 DEGs) of flg22-responsive genes were also regulated by Pep1 and only 21% (171 of 817 DEGs) were specific to flg22. We analysed how many genes respond in a cell type- and elicitor-specific manner. We pooled all elicitor-responsive DEGs to extract those genes that responded to one elicitor in at least one cell type and not to the other elicitor in any cell type. Splitting these genes up by cell types (Figure 4), we found that 89% (152 DEGs) of the flg22-specific DEGs are only expressed in one cell type with 96, 45 and 11 DEGs showing epidermis, cortex and pericycle-specific expression, respectively (Figure 4). In the epidermis and cortex, the majority of DEGs were induced whereas in the pericycle the majority of genes were suppressed (Supplemental Table 4, Supplemental Data Set 4). We performed a GO term analysis for each cell type and flg22-specific gene sets and found the flg22-specific epidermal expressed genes to be enriched in GO terms such as ‘regulation of proton transport’ and ‘response to nitrate’ and much less strongly enriched in immunity and defence associated terms (Supplemental Data Set 5). In the cortex the flg22-specific genes are enriched in the GO term ‘oligopeptide transport’ and ‘amide transport’, which matches the most strongly enriched GO terms from all the flg22-responsive genes specific to the cortex (from Figure 3E). This indicates that the enrichment of these peptide transport terms is a key cortex-responsive flg22-specific response.

**Figure 4.**
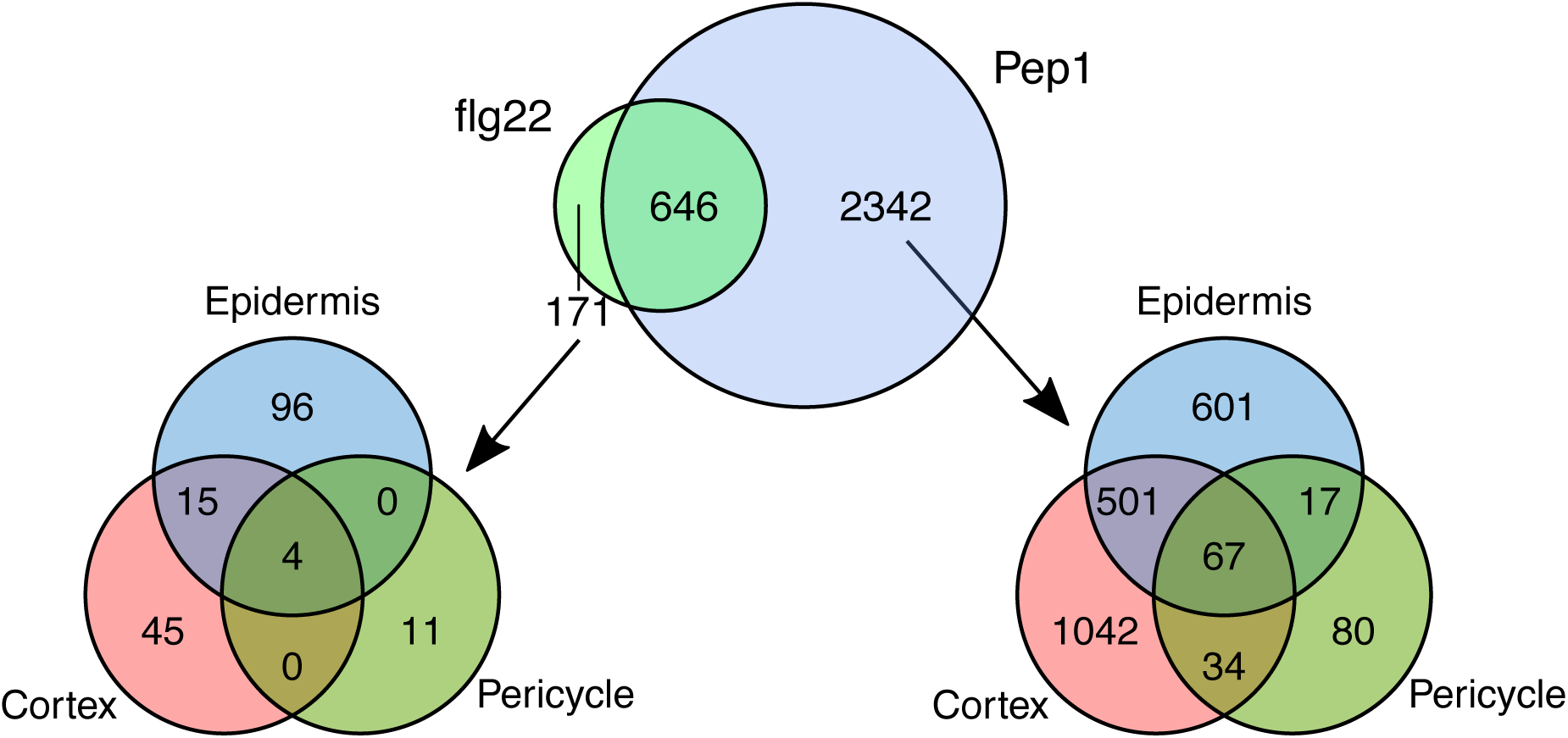
Differences between responses to flg22 and Pep1 in each cell type. Upper Venn diagram shows the overlap of flg22- and Pep1-responsive gene sets, aggregated across cell types. Lower Venn diagrams show the split by cell types for genes responding to flg22 in at least one cell type and not responding to Pep1 in any cell type (left) and vice versa (right).

For Pep1, 78% (2342 of 2988 DEGs) of the DEGs showed Pep1-specific expression out of which 74% (1723 DEGs) were expressed in only one root cell type with 601, 1,042 and 80 DEGs displaying epidermis, cortex and pericycle-specific expression, respectively (Figure 4). These cell type-specific DEGs had similar levels of up- and down-regulation across all three cell types (Supplemental Table 4, Supplemental Data Set 4). Consistent with the GO term enrichment of all Pep1-responsive genes, the Pep1-specific epidermal expressed genes were enriched in growth and hormones terms, particularly ‘ethylene biosynthetic process’, ‘SA biosynthetic process and ‘regulation of hormone levels’. In turn, the Pep1-specific cortex expressed genes were more strongly enriched in broad defence terms such as ‘response to biotic stimulus’ (Supplemental Data Set 5) encompassing genes such as *WRKY18, BIR1,* and *MYELOBLASTOSIS 122* (*MYB122*).

These data suggest firstly that the Pep1 response shows strong cell type specificity and that epidermis and cortex DEGs are indicative of contributing different functions to the Pep1 induced PTI. Moreover, these data indicate that, compared to Pep1, flg22 elicits a weaker response in root cells, and appears to be largely encompassed by the Pep1 response.

### Gene networks determining cell type immunity and cell identity are largely non-overlapping

Considering the substantial transcriptional changes in individual cell types upon immune elicitor treatments, we asked if immunity would affect the cell type identity. For each root cell type, identity is determined at the stem cell niche (Nakajima et al., 2001; Sabatini et al., 2003; Aida et al., 2004; Sarkar et al., 2007; Stahl et al., 2009). The maintenance of cell type identity, as defined by cell type-specific housekeeping functions, is of outstanding importance for overall root integrity and functionality (e.g. root growth), especially under stress (Iyer-Pascuzzi et al., 2011; Geng et al., 2013). Interestingly, immunity is known to halt root growth suggesting that housekeeping functions are over-ridden during root PTI (Gómez-Gómez et al., 1999; Jacobs et al., 2011). We therefore wanted to know if immunity affects cell type identity and with it the core of root tissue integrity and functionality.

We defined the cell type identity transcriptomes of unchallenged epidermis, cortex and pericycle cells. We identified 2,504 genes as specifically enriched in epidermis, 1,699 in cortex and 1,813 in pericycle (Figure 5A, Supplemental Data Set 6). These enriched datasets were confirmed to highly overlap (p<10^−6^) with published cell identity gene sets (Bargmann et al., 2013). Distinct GO terms were associated with each set of identity genes denoting the different functions of the three cell types (Figure 5B-D, Supplemental Data Set 6). The epidermis is enriched in cell division-associated GO terms (e.g. ‘cytokinesis by cell plate formation’) (Figure 5B), the cortex governs processes related to protein metabolism such as ‘Golgi organisation’ and ‘co-enzyme metabolic process’ (Figure 5C) and the pericycle is enriched in terms such as ‘S-glycoside biosynthetic process’ and ‘glucosinolate biosynthetic process’ indicative of secondary metabolism (Figure 5D).

**Figure 5.**
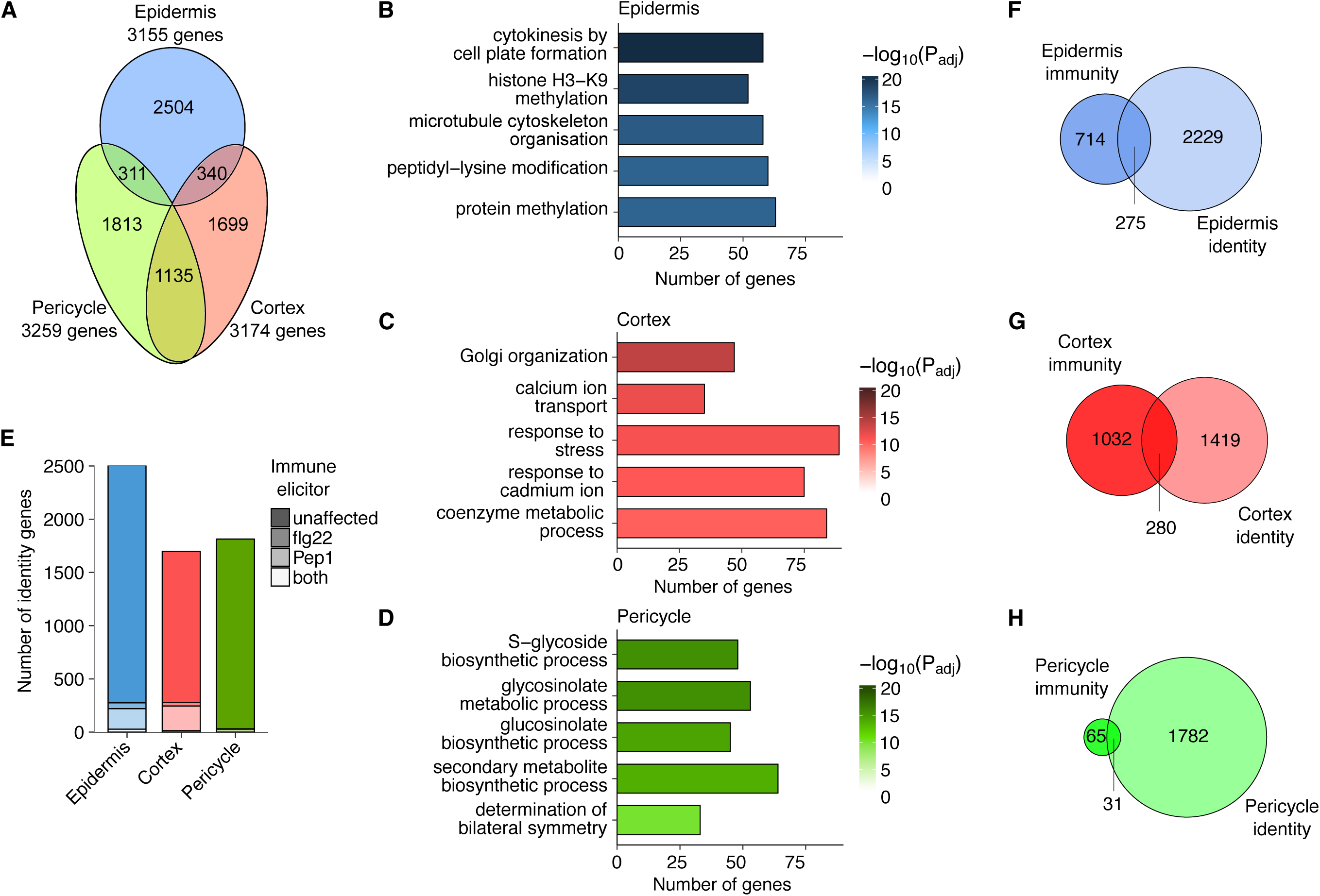
Immunity and cell identity. **(A)** Venn diagram showing numbers of up-regulated genes between cell types in untreated samples. For each cell type, genes that were up-regulated relative to at least one other cell type were identified. Non-overlapping areas termed ‘identity genes’. **(B)** Breakdown of immune elicitor response types for identity genes from (A). **(C), (D) and (E)** Top 5 enriched GO terms for identity genes. Only the 1,699 most significantly up-regulated genes from each cell type were used to make results more comparable across cell types. **(F), (G) and (H)** Venn diagrams showing the overlap between identity genes and genes DE in response to flg22 or Pep1 in only one cell type (termed immunity genes).

In order to answer the question of whether immunity affects cell type identity, we quantified the overlap between immunity genes and cell identity genes. We found that the regulation of only 11% of epidermis, 16% of cortex and 2% of pericycle-associated identity genes, respectively, was affected by either or both immune elicitor(s) (Figure 5E, Supplemental Data Set 7). On the other hand, the identity genes make up a larger proportion of the PTI response (to flg22 or Pep1) in each cell type. 28%, 21% and 32% of epidermis, cortex and pericycle PTI DEGs, respectively, are represented in the cell type identity gene sets (Figure 5F-H). These findings indicate the co-existence of cell identity and immunity-regulating networks in each cell type enabling highly cell type-dependent immunity responses without an overall effect on cell types. These findings are consistent with earlier studies, where cell identity was found to be unaffected by abiotic stress (Dinneny et al., 2008; Iyer-Pascuzzi et al., 2011; Geng et al., 2013).

### Distinct patterns of enriched combinatorial TF motifs in promoters of cell type-specific genes

We wanted to understand the regulation of cell identity and cell type-specific immunity networks. Therefore, we developed a new approach to identify enriched pairs of TF-binding motifs within the promoters of our gene sets of interest. Our analysis takes into account contributions of weak binding sites, multiplicity of binding sites for each factor, and accepts or rejects overlapping binding sites depending on the specificity of motifs in the region of the overlap (Figure 6A, see Materials and Methods). We opted to use a paired approach as one of the key features of transcriptional regulation in eukaryotes is combinatorial control by TFs. TF combinatorics have been shown to represent a key principle in regulating gene networks in *Arabidopsis* including immunity and hormones (Achard et al., 2009; Van de Velde et al., 2014; Lewis et al., 2015). In addition, by allowing motifs to overlap slightly, our analysis allowed us to consider TF complexes binding compound binding sites (Rodriguez-Martinez et al., 2017).

**Figure 6.**
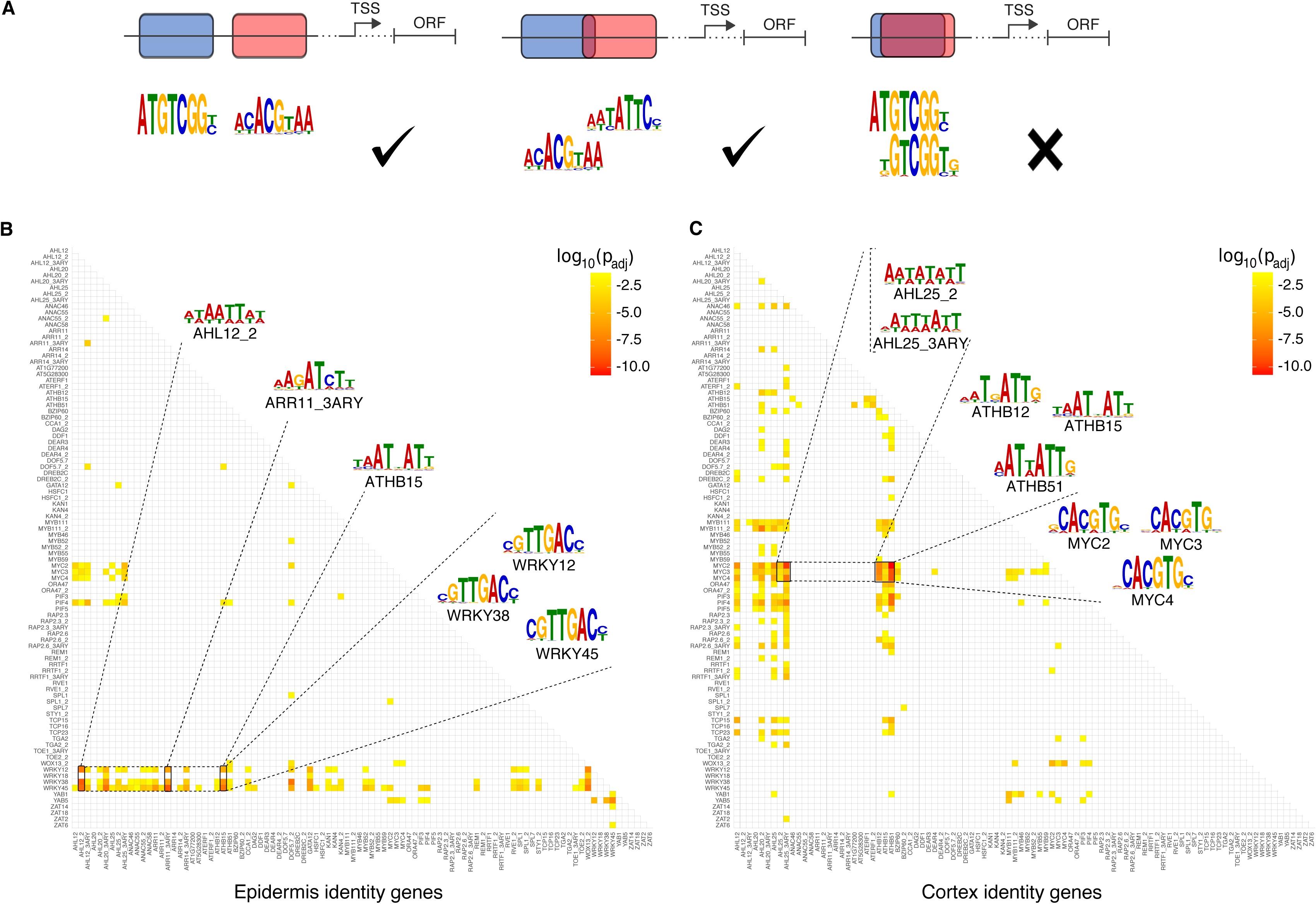
Different cell types show distinct patterns of paired-motif enrichment in cell type-specific genes. **(A)** Diagram illustrating the acceptance/rejection of overlaps in the paired motif enrichment analysis. **(B)** Heatmap representing p-values for enrichment of motif pairs in epidermis identity genes. WRKY motifs are found paired with a variety of motifs. **(C)** Heatmap representing p-values for enrichment of motif pairs in cortex identity genes. A pronounced feature is the pairing of AHL/ATHB motifs with MYC motifs. Motifs included and the order of motifs on both axes are the same as in (B).

First, we identified enriched motif pairs in our cell identity gene networks, using motifs from the motif database created by Franco-Zorrilla et al. (2014). To perform statiscally valid comparisons, we equalised datasets and used all 1,699 cortex-specific genes and the top 1,699 cell type enriched genes from the epidermis and pericycle (from Figure 5A). We found highly significant enrichment of a number of motif pairs in epidermis- and cortex-specific genes, but fewer and weaker signals for pericycle-specific genes (Figure 6B-C, Supplemental Figure 5, Supplemental Data Set 8). Enriched motifs covered the promoters of 1,002 epidermis (59%), 1,136 cortex (67%) and 337 pericycle (20%) cell identity genes. Comparing the epidermis and cortex, the pattern of enriched motif combinations was highly distinctive with a large number of motif pairs highly significant in one cell type and not statistically significant in the other.

Within the promoters of epidermis identity gene motifs three different WRKY TFs (WRKY12/38/45) were found to pair with a wide range of motifs (Figure 6B). In particular, WRKYs were enriched with motifs for AT-HOOK MOTIF CONTAINING NUCLEAR LOCALIZED (AHLs) (p<10^−8^, Bonferroni-corrected p-value corresponding to enrichment score of AHL12_2 and WRKY45 motif pairs), ARABIDOPSIS RESPONSE REGULATORs (ARRs) (p<10^−9,^ Bonferroni-corrected p-value corresponding to enrichment score of ARR11_3ARY and WRKY45 motif pairs), and ARABIDOPSIS THALIANA HOMEOBOX (ATHBs) TFs (p<10^−7,^ Bonferroni-corrected p-value corresponding to enrichment score of ATHB15 and WRKY45 motif pairs), accounting for 26% of all epidermis identity genes (442 genes of 1,699 genes). WRKYs represent a large family of *Arabidopsis* TFs (>70 members) with known regulatory functions in plant innate immunity, abiotic stress and developmental processes (Pandey and Somssich, 2009; Rushton et al., 2010). In turn, AHL and ARR TFs have distinct roles in growth and development with ARRs of particular importance in cytokinin signalling (Matsushita et al., 2007; Ishida et al., 2008; Kushwah et al., 2011; Hur et al., 2015). In the cortex, in turn, gene promoters showed enriched pairing of MYC and PHYTCHROME-INTERACTING FACTOR (PIF) TF binding motifs with AHL and ATHB TF binding motifs (Figure 6C). These are further examples of the pairing of stress and development-related TFs. MYC2-4 motifs identified in our analyses are well known integrators of plant immune signalling (Fernández-Calvo et al., 2011; Schweizer et al., 2013) whereas PIFs and ATHBs take key roles in adapting growth according to environmental conditions such as shade avoidance or photocontrol (Prigge et al., 2005; Leivar and Quail, 2011). We noted that the overlap in enriched motif pairs between epidermis and cortex was very small, mostly limited to MYCs showing pairing with AHL TF motifs for both tissues (Figure 6B-C). Overall, our paired motif analyses detected co-occurring motif pairs with a predominant gene network regulatory role of WRKY, AHL and ARR combinations in the epidermis and MYC, PIF, AHL and ATHB TF combinations in the cortex.

### WRKYs cooperate with developmental TFs to regulate cell type-specificity of flg22 immunity

We conducted a paired motif enrichment analysis to understand if the cell type-specific rewiring of gene networks upon flg22 is reflected by a distinct enrichment in promoter motif pairs. Therefore, we tested the promoters of genes specifically induced in the epidermis or cortex in response to flg22. To make the enrichment scores across flg22-treated cell types directly comparable we equalised gene set sizes by taking all of the cortex-specifically up-regulated genes (120 genes, from Figure 3B) and the top 120 most significantly up-regulated, epidermis-specific genes. The analysis revealed an enrichment of highly specific motif pairs in the promoters of 78 (out of 120) epidermis and 88 (out of 120) cortex genes up-regulated by flg22. Overall, the promoters of flg22-responsive DEGs were particularly enriched in WRKY motifs for WRKY12, WRKY18, WRKY38 and WRKY45. In the epidermis, these WRKY motifs showed particularly enriched pairing with KAN1, KAN4 and KAN4_2 motifs (p<10^−7^; Bonferroni-corrected p-value corresponding to enrichment score of KAN4 and WRKY45 motif pairs), which bind the KANADI family of TFs (Figure 7A). KANADIs have been shown to act as a negative regulator in embryo development (McAbee et al., 2006), root development (Hawker and Bowman, 2004) and vascular tissue formation (Ilegems et al., 2010).

**Figure 7.**
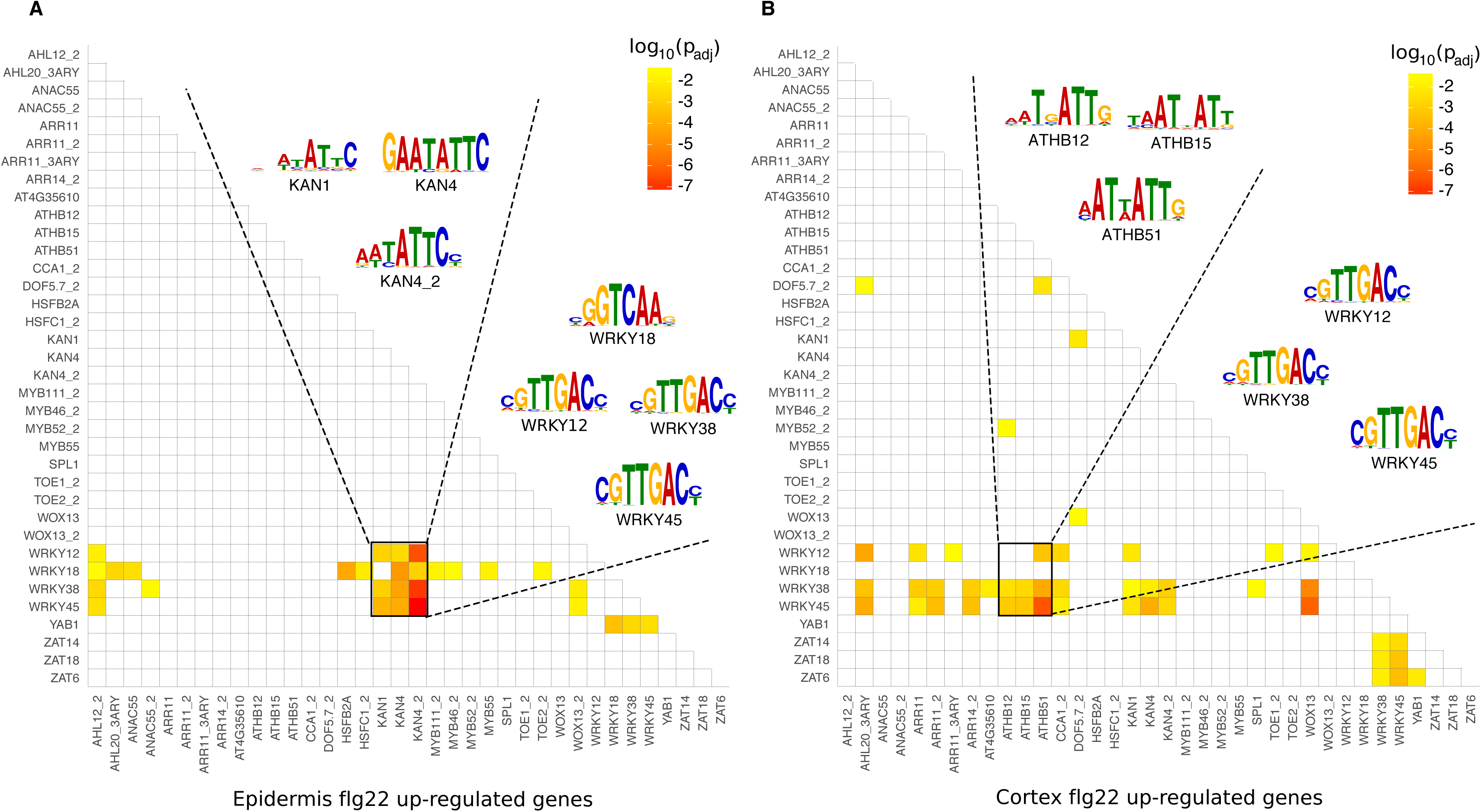
Paired motif enrichment of cell type-specific flg22-response genes. **(A) and (B)** Heatmap representing p-values for enrichment of motif pairs in promoters of genes up-regulated in response to flg22 specifically in the epidermis (A) and the cortex (B). Motifs included and the order of motifs on both axes are the same in (A) and (B).

We asked if the expression pattern of these TFs was matched with the pattern of enrichment of their target sites. KAN1, KAN2 and KAN4 were specifically enriched in the epidermis dataset and not in either of cortex and pericycle, consistent with the gene expression database ePlant (Waese et al., 2017). KAN/WRKY motif pairs were predominant and present in the promoters of 42 of the 120 flg22-induced epidermis genes and this gene set included core immune signalling receptor-like kinases from the *CRK* and *WAKL* families. The combination of motif enrichment and expression patterns suggests that the KAN/WRKY paired motifs are an epidermis-specific elicitor signalling mechanism (Figure 7A, Supplemental Data Set 9). In the cortex, this enrichment of WRKY/KAN motif pairs was weaker and only partially seen (p<0.005, Bonferroni-corrected p-value). Instead, motif enrichment patterns were very different between epidermis- and cortex-specific gene sets. WRKYs (especially WRKY38 and WRKY45) showed enriched pairing with ARR (ARR11/14), ATHB (ATHB12/15/51) and ZINC FINGER OF ARABIDOPSIS THALIANA (ZAT) motifs (ZAT6/14/18) (Figure 7B) in cortex cells. ZATs appear to have a general role in mediating abiotic stress tolerance (e.g. cold, drought) (Yin et al., 2017), while ZAT14 integrates phosphate starvation responses (Devaiah et al., 2007). Interestingly, WRKY18 showed a clear cell type-specific pattern. While WRKY18 showed enriched pairing with a number of motifs (AHL12_2, AHL20_3ARY, ANAC55, HSFB2A, HSFC1_2, KAN4, KAN4_2, MYB111_2, MYB46_2, MYB55, TOE2_2, YAB1) in the epidermis, it did not pair with any motif in the cortex; a pattern very reminiscent to the overall WRKY18 absence of paired enrichment in promoters of epidermis cell identity genes (Figure 6B).

Comparing the results for the flg22 and cell identity gene sets revealed some overlaps in motif pair enrichment, e.g. for WRKY/AHL and WRKY/KAN motifs, which were only observed for epidermis data.

### Context dependent linkage of immunity and cell identity networks

We were interested to see if our paired motif enrichment analyses can further explain overlaps between cell type-specific immunity and identity networks. First, we performed paired enrichment analyses with cell type-specific DEGs after Pep1 treatment. Again, we equalised gene set sizes by including all epidermis-specifically Pep1-induced and Pep1-suppressed genes (370 and 358 genes, respectively; Figure 3C) as well as the 370 and 358 most significantly Pep1-induced and -suppressed, cortex-specific genes, respectively. The analyses identified paired motif enrichment in the promoters of 162 and 235 out of 370 epidermis- or cortex-induced genes, respectively, and 179 and 181 out of 358 epidermis- or cortex-suppressed genes, respectively (Figure 8, Supplemental Data Set 10). For Pep1-induced genes in the epidermis, we identified KAN4 to show enriched pairing with WRKY12/38/45 (found in 48 out of 370 tested promoters; Figure 8A). In turn, the analysis of epidermis-specific Pep1-suppressed genes revealed AHLs (AHL20_2/AHL25) to pair with MYC2/3/4 and PIF4 (found in 61 out of 358 promoters; Figure 8C). For cortex-, Pep1-induced genes, most dominant was the pairing of MYC (MYC2/3/4) with WRKY TFs (WRKY12/38/45), a pairing that was found in 47 out of 370 promoters of cortex-specifically Pep1-induced genes (Figure 8B). Finally, we detected enriched pairing of MYB TF motifs (MYB111/MYB111_2/MYB46) particularly with ATHBs (ATHB12/15/51), found in 48 promoters of 358 cortex-specifically Pep1-suppressed DEGs (Figure 8D). MYB46 and MYB111 have been reported to be involved in flavonol glycoside metabolism (Stracke et al., 2010) and secondary cell wall synthesis, respectively (Zhong et al., 2007; Ko et al., 2009). Interestingly, MYC TF motifs were enriched in promoters of Pep1-induced genes in the cortex (paired with WRKYs) and Pep1-suppressed genes in the epidermis (paired with AHLs). This finding indicates the efficiency of paired motif enrichment analysis and its potential in interpreting previous reports. MYC TFs have been implicated in Pep1-mediated signalling in particular as Peps specifically induce the MYC2-dependent branch of jasmonic acid (JA)-responsive signalling (Bartels and Boller, 2015). Additionally, MYC2 has been shown to act as both an activator and a repressor in JA-mediated gene expression (Dombrecht et al., 2007). Comparing cell type-specific identity and immunity networks (using flg22/Pep1-induced genes) based on our paired motif enrichment analyses, we observed significant patterns (Figure 9). In the epidermis, WRKY12/18/38/45 connects identity with immunity networks by pairing with KAN1/4 (dominating epidermis immunity networks) and with AHL12/ARR11/ATHB15 (dominating epidermis identity networks), respectively (Figure 9). In turn, in the cortex, MYC2/3/4, PIF3/4/5 and ATHB12/15/51 ties both networks by pairing with WRKY12/38/45 (dominating cortex immunity networks) and with AHL12/20/25 (dominating cortex identity networks), respectively (Figure 9). Our paired motif enrichment analyses suggest that cell identity integrates with and determines root immunity by specific TF pairs in individual cell types. In summary, the patterns of motif pairing that are statistically linked with our cell type-resolved expression data provide distinctiveness across cell types, across treatments, and in comparisons of gene induction vs. gene suppression. Moreover, it allows the identification of potential regulatory mechanisms to connect cell identity with immunity networks.

**Figure 8.**
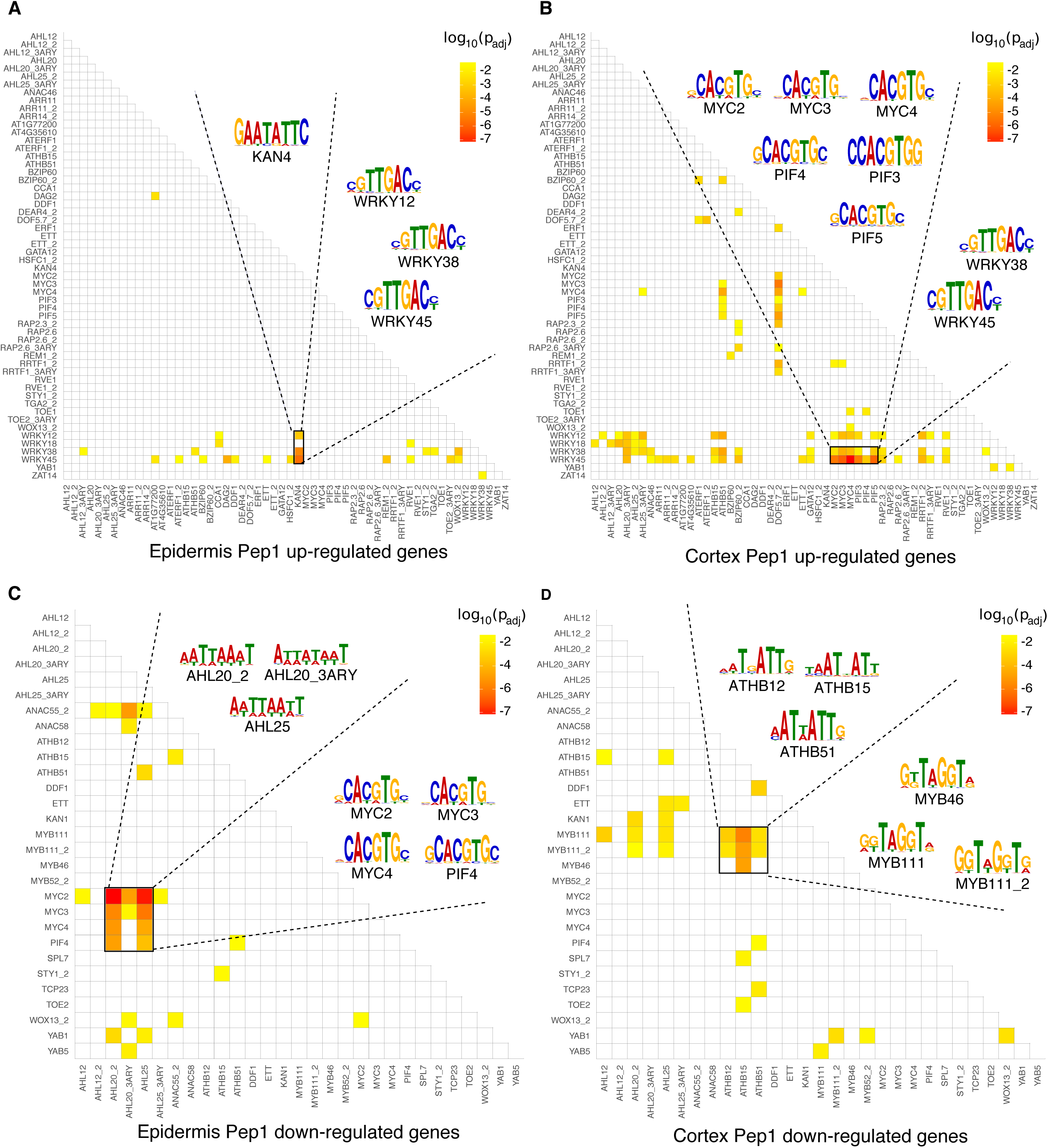
Paired motif enrichment of cell type-specific Pep1-response genes. **(A) and (B)** Heatmap representing p-values for enrichment of motif pairs in promoters of genes up-regulated in response to Pep1 specifically in the epidermis (A) and the cortex (B). Motifs included and the order of motifs on both axes are the same in (A) and (B). **(C) and (D)** Heatmap representing p-values for enrichment of motif pairs in promoters of genes down-regulated in response to Pep1 specifically in the epidermis (C) and the cortex (D). Motifs included and the order of motifs on both axes are the same in (C) and (D).

## Discussion

In this study we have analysed the immune responsiveness of three root cell types as a first step to decipher the coordination of immunity in a complex tissue. We studied epidermis and cortex cells, which build the outer frontier to the rhizosphere, as well as the pericycle as outer frontier of the inner root vasculature. Based on their location, these cell types have an important role in root immunity. Our analyses reveal that different root cell types are not only highly immune responsive but respond very differently to the immune elicitors flg22 and Pep1. By comparing the responses to flg22 and Pep1, we were able to elucidate specific gene networks for each of the three root cell types. Both elicitors activate different networks in each cell type demonstrating a remarkable complexity of immunity in roots. Although it may require a higher degree of coordination (e.g. numerous regulatory and signalling proteins), this cell type-specific gene network regulation may add the robustness and flexibility needed for a root system to adapt to constantly changing environmental stimuli. Consistent with this, recent studies suggested qualitative differences in the immune competences of different root cell types (Beck et al., 2014; Wyrsch et al., 2015; Poncini et al., 2017). By expressing the flg22 receptor FLS2 in a cell type- and root development-specific manner, Wyrsch et al. (2015) observed flg22 responsiveness in cell types across different root development zones. By analysing defined PTI responses (e.g. MAPK phosphorylation, immunity marker gene expression) their study suggested different contributions of cell types to PTI. Similarly, studies with *Arabidopsis* roots exposed to abiotic stress indicated cell type-specificity in stress integration. Irrespective of the nature of the abiotic stress (e.g. salt stress, iron starvation, nitrogen depletion) each cell type responded differently and in a highly coordinated manner to maintain root functionality under stress (Iyer-Pascuzzi et al., 2011). For instance, roots showed a transient root growth inhibition phenotype under salt stress that coincided with the cell type-specific rewiring of hormone signalling to reconfigure root growth-regulating networks under this environment (Geng et al., 2013). Such plasticity in root growth and development appears to be fundamental as it has been observed under nitrogen depletion and was shown to drive lateral root development (Gifford et al., 2008; Walker et al., 2017). It further underlines the hierarchy of root (cell) function with the maintenance of growth and development as top priorities under fluctuating environments. Compromising cell identity would jeopardise root tissue function and hence, plant fitness and survival. Accordingly, our study demonstrates that root cell types keep their identity under biotic stress. We observed that housekeeping and stress-responsive gene networks co-exist in each root cell type, and the overlap could be of importance to overall root plasticity (Figure 9). Our data support the concept that cell identity underpins transcriptional reprogramming leading to exquisite cell specificity in response to signal perception, even modulating outputs from strong elicitors such as MAMPs. In line with previous studies on abiotic stress integration (Dinneny et al., 2008; Geng et al., 2013), our data suggest that cell identity networks underpin the cell type-specific immune responses. Linking immunity to cell identity networks would guarantee cell type-specific regulation of immune responses according to the functional competence of each cell type. Such a co-regulatory model would likely be applicable to all environmental stresses.

**Figure 9.**
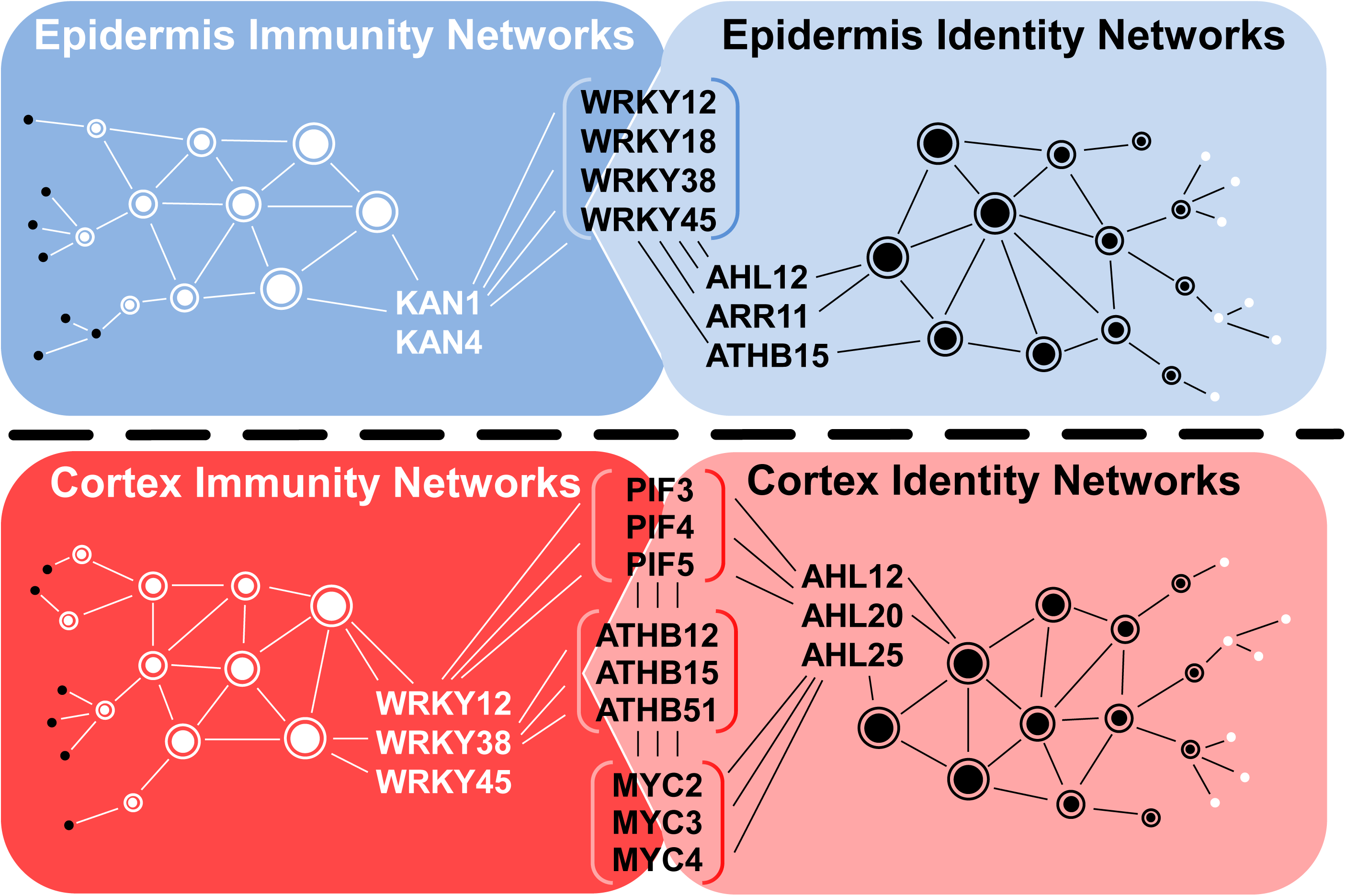
Model for the connection between cell type-specific identity and immunity networks. Connection of epidermis and cortex-specific cell identity networks with immunity networks via transcriptions factors (in brackets) that are part of respective immunity and identity networks.

Our high resolution, combinatorial promoter analysis provided new and important insights into how immunity and cell identity networks are coordinated. It enabled us to explain differences in gene regulation between cell types and treatments with statistical confidence. Importantly, such a combinatorial analysis allowed us to consider cooperative binding of different TFs, which can enhance the flexibility of gene regulation under changing environments (Van de Velde et al., 2014). The inclusion of two promoter motif binding sites was sufficient to explain the regulation patterns of 44-73% of DEGs per cell type and treatment. We found that pairing of TF motifs differed dependent on the cell type suggesting that different TF families act together to create the highly cell type-specific networks that we observed, and these can be further tuned by specific combinations of TF family members. We noted a striking pattern in the pairing of promoter motifs for known stress-regulatory TF families with motifs for developmental TFs in each cell type-specific gene network. Moreover, certain TF combinations prevailed in specific cell types in a treatment-specific manner. For instance, WRKY TFs might have a more prominent function in regulating epidermis-specific networks together with specific developmental TFs, with WRKY/AHL, WRKY/ATHB and WRKY/ARR paired motifs regulating cell identity networks and KAN/WRKY paired motifs regulating cell immunity networks. In turn, cortex function relies on MYC, ATHB, and PIF TFs in combination with WRKY TFs to regulate cortex-specific immunity and with AHLs to regulate cortex identity-specific gene networks. It will be interesting to explore in future studies to what extent this combination of stress and developmental TFs contribute to stress integration and, especially to growth regulation under immunity. Furthermore, our paired motif analyses allowed us to distinguish between elicitor-specific networks in each cell type. These differences between PTI elicitors might reflect the life strategies of pathogens with flg22-induced PTI evolved to defend against bacteria. Pep1 in turn is an intrinsic PTI elicitor that is activated by different hormones and, hence, might trigger the full array of all immune responses against a larger variety of pathogens.

The order of complexity of cell type-specific immunity requires a tight coordination. Our data strongly suggest that TF combinations mediate this complexity by cell type-specific gene regulation. Our study provides the first insight into cell type-specific immunity networks, and combinatorial TF motifs associated with specific responses and cell types. Such analyses might further help in interpreting GO term enrichment and potentially redefining GO terms as we could assign genes based on regulatory promoter motifs in addition to gene expression patterns. A striking difference was observed for the context-dependent association of MYC motifs in Pep1 responses. For promoters of genes specifically up-regulated in cortex cells, MYC2/3/4 were found to pair with WRKYs, with MYC4 showing the strongest statistical link. In contrast, MYC2/3/4 were found to be paired with AHL20/25 in promoters of down-regulated genes in the epidermis, with MYC2 showing the strongest association. Neither up-regulation in epidermis nor down-regulation in cortex were found to be linked to either of these motif combinations. Thus our analyses reveal potential exquisite multi-functionality of specific TF family members whereby they can act with conserved but heterogeneous partners in different cell types to effect contrasting transcriptional outputs. Our findings highlight how context dependency of regulatory function may reduce perceived redundancy among regulatory factors recognising highly similar sites if investigated in isolation.

## Materials and Methods

### Plant materials, growth conditions and treatment

Seeds of *A. thaliana* ecotype Col-0 were obtained from the Nottingham *Arabidopsis* Stock Center. Seeds of the *pepr1-2 pepr2-2* mutant were kindly provided by Y. Yamaguchi, Osaka Prefecture University, Osaka, Japan. Marker lines for the three cell types *pGL2:GFP* (epidermis), *pCORTEX:GFP* (cortex), and *E3754* (pericycle) were obtained from Miriam Gifford, University of Warwick, UK. Plants were grown on vertical squared Petri dishes on ATS medium (Lincoln, 1990) without sucrose and supplemented with 4.5 g l^-1^ Gelrite (Duchefa Biochemie) in a 22°C day/18°C night cycle (8 h light) at 120 μmol m^-2^s^-1^. For experiments with *S. indica*, roots of 9-10-days-old plants were inoculated with 1 ml of a 500,000 chlamydospores ml^-1^ spore suspension per squared Petri dish. Control plants were treated with H_2_O_0.02% Tween20_ (mock). If not stated otherwise, plants were treated on plates with 1 ml per plate of 1 μM solutions of flg22 or Pep1 or H_2_O as control. For all experiments, flg22 and Pep1 peptides were used as described (Gómez-Gómez et al., 1999; Krol et al., 2010). All data is based on at least three biological experiments.

### Measurement of plant growth inhibition

For seedling growth inhibition assays, plants were grown on squared Petri dishes for 10 d before inoculation with *S. indica* or mock treatment. After three days, plants were transferred into round Petri dishes containing liquid ATS medium supplemented with 1 μM flg22, 1 μM Pep1 or water (control). Eleven days later, plant fresh weights were determined. In each of the three biological experiments, 12 plants per line and treatment were evaluated.

### Measurement of ROS burst

Roots of 2-weeks-old plants were grown on solid half-strength MS medium and treated with 1 μM flg22, 1 μM Pep1 or mock at 3 days after inoculation with *S. indica* or mock treatment. For ROS burst quantification, roots were cut in 1-cm-long pieces (10 mg per assay) and transferred to a luminol-based assay as described (Gómez-Gómez et al., 1999). Data were analysed by Student’s *t* test.

### MAPK protein and phosphorylation assay

Roots of 21-days-old *Arabidopsis* seedlings were harvested into liquid nitrogen 10, 30 and 60 mins after immune elicitor treatment. Total protein was extracted after grinding and homogenising material in protein extraction buffer containing Tris-HCl (pH 7.8) 25 mM NaCl, 75 mM EGTA, 15 mM MgCl_2_, 10 mM Tween 20 0.1% (v/v), PMSF, 0,5 mM leupeptin, 10 μg μl^-1^ aprotinin, 10 μg μl^-1^ glycerophosphate, 15 mM Tris, 15 mM NaF, 1 mM Na_3_VO_4_, 0.5 mM DTT. 30-40 μg of total protein extract were subjected to SDS-PAGE. After transfer to nitrocellulose membrane, proteins were incubated with monoclonal mouse anti-Phospho-p44/42 MAPK (Erk1/2, Thr202/Tyr204, 1:1,000 dilution) antibody (Cell Signalling Technology), anti-MPK6 (1:10,000), anti-MPK3 (1:5,000) and anti-MPK4 (1:5,000) antibodies (all Sigma Aldrich). Antibodies were diluted in 5% BSA in TBS-T. After replacing primary with secondary anti-rabbit IgG HRP-conjugated antibody (1:10,000, Sigma Aldrich), the samples were incubated for two h before signal detection using a Femto-ECL kit (Pierce) and Amersham Hyperfilm (GE Healthcare).

### Gene expression analysis using qRT-PCR

For gene expression analyses in whole roots, root material was harvested 2 and 24 h after flg22, Pep1 or control treatment. TRIzol (Invitrogen) was used to extract total RNA. After DNAse treatment, RNA was reverse transcribed into cDNA using the qScript cDNA synthesis kit (Quanta Biosciences Inc.). We used 10 ng of cDNA as template in quantitative real-time PCR (qRT-PCR) reactions using the SYBR Green JumpStart Taq ReadyMix (Sigma-Aldrich) in a Stratagene Mx3005P Real-time PCR Detection System (Agilent Technologies) following a standard protocol. The 2^-ΔCt^ method (Schmittgen and Livak, 2008) was used to determine differential expression of marker genes for immunity activation (for primer sequences see Supplemental Table 5).

### Quantification of *S. indica* colonisation by qRT-PCR

Genomic DNA was isolated from roots using the Plant DNeasy Kit (Qiagen). 40 ng of genomic DNA served as template in quantitative real-time PCR (qRT-PCR) reactions using the SYBR Green JumpStart Taq ReadyMix (Sigma-Aldrich) in a Stratagene Mx3005P Real-time PCR Detection System (Agilent Technologies) following a standard protocol. The 2^-ΔCt^ method (Schmittgen and Livak, 2008) was used to determine fungal colonization by subtracting the raw cycle threshold values of *S. indica* internal transcribed spacer (ITS) from those of *Arabidopsis UBQ5* (for primer sequences, see Supplemental Table 5). Data were analysed by Student’s *t* test.

### Fluorescence-activated cell sorting (FACS)

For FACS experiments, plants were grown for 12 days on squared ATS plates, and then treated with 1 ml per plate of 1 μM solutions of flg22 or Pep1 peptide or H_2_O as control for 1 h. Whole roots were cut into pieces and then incubated in protoplast solution (1.5% cellulase R10 (Duchefa Biochemie), 1.2% cellulase RS (Duchefa Biochemie), 0.2% macerozyme R10 (Duchefa Biochemie), and 0.12% pectinase (Sigma Aldrich), in 600 mM mannitol, 2 mM MES hydrate, 10 mM KCl, 2 mM CaCl_2_, 2 mM MgCl_2_, and 0.1% BSA, pH 5.7) (Walker et al., 2017) for 1 h. Protoplasts were filtered through 70 and then 40 μm cell strainers, centrifuged at 300 g for 3 mins, resuspended in protoplast solution lacking cell wall-degrading enzymes and subjected to FACS. Three independent biological experiments were carried out for each marker line. GFP-expressing protoplasts were collected by using BD Influx cell sorter (BD Biosciences), following previously published protocols (Birnbaum, 2003; Gifford et al., 2008; Grønlund et al., 2012)The cell sorter was equipped with a 100 μm nozzle and BD FACSFlow^™^ (BD Biosciences) was used as sheath fluid. BD^™^ Accudrop Fluorescent Beads (BD Biosciences) were used prior to each experiment to optimize sorting settings. A pressure of 20 psi (sheath) and 21 – 21.5 psi (sample) was applied during experiments. Drop frequency was set to 39.2 kHz, and event rate was generally kept < 4000 events s^-1^. GFP-expressing protoplasts were identified using a 488 nm argon laser, plotting the outcome of a 580/30 bandpass filter vs. a 530/40 bandpass filter, to differentiate between green fluorescence and autofluorescence. Different cell populations were collected for microscopy in pre-experiments to determine the presence of GFP-expressing protoplasts. As previously published (Grønlund et al., 2012), these protoplasts were present in the high 530 nm / low 580 nm population. Sorting gates were set conservatively in following experiments based on these observations (Supplemental Figure 6). For RNA-extraction, GFP-expressing protoplasts were sorted into Qiagen RLT lysis buffer containing 1% (v:v) β-mercaptoethanol, mixed, and immediately frozen at −80°C. At least 10,000 GFP-expressing protoplasts were sorted per experiment and treatment condition. Sorting times were kept below 25 mins.

### RNA isolation, RNA-seq library construction and sequencing

Total RNA was extracted using the Qiagen RNeasy Plant Mini Kit including on-column DNase treatment with the Qiagen DNase kit. The 6000 Pico Kit (Agilent Technologies) was used to check quantity and quality of the RNA on a Bioanalyzer 2100 (Agilent Technologies). Preparation of amplified cDNA from total RNA and library construction were done with the Ovation^®^ RNA-Seq System V2 and Ovation^®^ Ultralow Library Systems Kit (NuGEN Technologies), respectively, following standard protocols. Sequencing was carried out by the High-Throughput Genomics Group at the Wellcome Trust Centre for Human Genetics on an Illumina HiSeq2500 System.

### RNA-seq quality control and read mapping

For each sample, read quality was evaluated using the FastQC software (Andrews, 2010). The paired-end libraries (2 × 100 bp reads) were mapped to the *A. thaliana* TAIR10 genome using Tophat2 (Kim et al., 2013) (default parameters –library type unstranded (Langmead and Salzberg, 2012)). The reads mapping to exons were counted using HTSeq-count (Anders et al., 2015) (settings: -f bam -s no -i Parent -t mRNA) with a GTF annotation file from the TAIR website (www.arabidopsis.org, downloaded October 2016). On average 48.1 percent of reads uniquely mapped to exons (full details in Supplemental Table 1). The quality of read mapping was assessed using the Integrative Genomics Viewer (IGV) (Robinson et al., 2011). The quality of replicates was assessed by plotting read counts of samples against one another and assessing the dispersion and presence of any artefacts between samples. Due to preferential amplification in some samples, reads corresponding to rRNA and ribosomal proteins had to be removed for subsequent analyses (Supplemental Data Set 11). The mitochondrial and plastid chromosomes were also removed as this work focuses on nuclear-encoded genes. Principal component analysis was calculated using the function prcomp in R and visualised in MATLAB.

### Differential gene expression and functional analysis

Differentially expressed genes (DEGs) were identified using DESeq2 (Love et al., 2014) using the paired replicate approach and filtered using an adjusted p-value (FDR) of 0.05. Immunity genes were determined by comparing treatments and mock treatments within each cell type. Cell type identity genes were identified using mock-treated samples only. The fit of the DESeq2 model to our data was tested by plotting replicates against one another and overlaying the read counts of DEGs (Supplemental Figure 4). Subset analysis to determine cell type and treatment-exclusive genes was performed in R using inbuilt set functions and the “VennDiagram” (Chen and Boutros, 2011) package. Proportional visualisations of 3-set Venn diagrams were created using “EulerAPE” (Micallef and Rodgers, 2014), non-proportional Venns were created using the venn function from the R packaged “gplots”, 6 set Venn diagram was created using interactive Venn tool by (Bardou et al., 2014). Gene Ontology (GO) enrichment analysis was performed using the R package “GOStats” (Falcon and Gentleman, 2007) with an additional Benjamini-Hochberg multiple testing correction applied. To make the enrichment scores across cell types directly comparable we equalised gene set sizes by taking the top *K* genes from the larger dataset where *K* is the size of the smaller dataset.

### Paired motif enrichment analysis

113 motifs (in the form of letter-probability matrices) were obtained from microarray studies performed by Franco-Zorrilla et al., 2014. These motifs characterize the target sequence specificity of 63 plant TFs representing 25 families. Promoter regions corresponding to 1,000 bp upstream from the transcription start site were collected from the TAIR10 database (www.arabidopsis.org) for all nuclear genes (including transposable element genes and excluding genes on the mitochondrial or plastid chromosomes) in the *Arabidopsis* genome. For each motif and each promoter, the sequence was scanned for occurrences of the motif using FIMO (Grant et al., 2011) which assigns a probability score to each potential hit. In order to determine the number of hits to consider and to compute an overall score for motif presence in a promoter, we computed the geometric mean *p* of the top *k* FIMO probability scores for non-overlapping hits and computed the binomial probability of observing at least *k* hits of probability *p* in a 1 kb promoter. The value of *k* minimising the binomial probability was taken to indicate the most likely number of binding sites and the *k* hits were recorded for subsequent analysis (1 ≤ *k* ≤ 5). For each motif, the promoters were ordered by increasing binomial probability and the top *n* = 5,000 promoters were considered as containing the motif. The parameter *n* was chosen for high sensitivity (rather than specificity) as stringency is introduced when the pairing of motifs is considered. The binomial probability of the *n*-th promoter was recorded for each motif as a threshold. For each pair of motifs and for each promoter containing both motifs, overlaps of recorded motif hits were identified and the information content (IC) of the overlap (based on motifs) was calculated. If the IC of the overlap for either motif exceeded 4 (indicating highly conserved bases are part of the overlap), then these hits were removed and the binomial probability re-calculated for the remaining hits. If the re-calculated scores were still below the recorded motif-specific threshold, then the two motifs were considered as co-localised in the promoter. Finally, gene sets of interest were tested for enrichment of paired motifs using a pairwise hypergeometric test based on the MATLAB function proposed by Meng et al., (2009). Hypergeometric p-values were corrected for the number of motif pairs using “local” Bonferroni (calculating the correction for each gene set separately). Corrected p-values<0.05 are considered significant. For each comparison of results made between conditions, the gene sets tested were of equal size to make p-values comparable. To this end, gene set sizes were equalised by taking the top *K* genes from the larger gene set, where *K* is the size of the smaller gene set.

## Supplemental Data

**Supplemental Figure 1.** The ability of *Serendipita indica* to suppress flg22 but not Pep1-triggered immune responses in roots.

**Supplemental Figure 2.** Assessment of RNA-seq data quality.

**Supplemental Figure 3.** 6-set Venn diagram detailing all overlaps between sets of DEGs for the two treatments in three cell types.

**Supplemental Figure 4.** Visual inspection of differential gene expression.

**Supplemental Figure 5.** Heatmap representing p-values for enrichment of motif pairs in pericycle identity genes.

**Supplemental Figure 6.** Representative FACS dot plots of the output from the 580/30 nm vs. the 530/40 nm bandpass filters.

**Supplemental Table 1.** Alignment statistics table based on output from htseq-count.

**Supplemental Table 2.** Numbers of DEGs responding to flg22 and Pep1 in three root cell types.

**Supplemental Table 3.** Genes differentially expressed in all three cell types in response to both flg22 and Pep1.

**Supplemental Table 4.** Numbers of DEGs responding specifically to either flg22 or Pep1 in one or more cell types.

**Supplemental Table 5.** Primer sequences

**Supplemental Data Set 1.** Lists of all differentially expressed genes in response to flg22 and Pep1 from DESeq2 output.

**Supplemental Data Set 2.** Lists of cell type-specific DEGs following flg22 or Pep1 treatment.

**Supplemental Data Set 3.** Lists of cell type-specific GO terms after flg22 and Pep1.

**Supplemental Data Set 4.** Lists of flg22- and Pep1-specific DEGs per cell type.

**Supplemental Data Set 5.** Lists of flg22- and Pep1-specific GO terms per cell type.

**Supplemental Data Set 6.** Lists of cell identity genes and enriched GO terms.

**Supplemental Data Set 7.** Lists of PTI genes within cell type-specific cell identity genes.

**Supplemental Data Set 8.** Paired motif analysis results for cell identity genes.

**Supplemental Data Set 9.** Paired motif analysis results for flg22 up-regulated genes.

**Supplemental Data Set 10.** Paired motif analysis results for Pep1 up- and down-regulated genes.

**Supplemental Data Set 11.** Genes omitted from the analysis. See Materials and Methods.

## Author Contributions

M.R., K.H.K., S.O. and P.S. designed research; C.R., M.R., S.H., D.J.J. and R.E. performed research; C.R. and E.E. contributed new computational tools, all authors analysed data and were involved in writing the paper.

## Acknowledgements

We thank Murray Grant for helpful comments on the manuscript, Jeanette Selby and Lesley Ward at the Genome Research Facility (University of Warwick, School of Life Sciences) for their help in RNA-seq library preparations, Christina Neumann and Rebekka Schmitt (Institute of Phytopathology, Justus Liebig University Giessen) for help with PTI assays, Nicholas Provart and Asher Pasha (Department of Cell and Systems Biology / Centre for the Analysis of Genome Evolution and Function, University of Toronto) for help with ePlant data deposition and view creations. We thank the High-Throughput Genomics Group at the Wellcome Trust Centre for Human Genetics for the generation of sequencing data. This work was funded by research grants from Biotechnological and Biological Research Council (BBSRC) / Engineering and Physical Sciences Research Council Grant (EPSRC) of the United Kingdom and the Deutsche Forschungsgemeinschaft (DFG), with grant IDs: BB/M017982/1 (PS), SCHA1444/3-3, SCHA1444/5-2 (both PS). CR was funded by the Biotechnology and Biological Sciences Research Council through MIBTP.

---

**Supplemental Figure 1.**
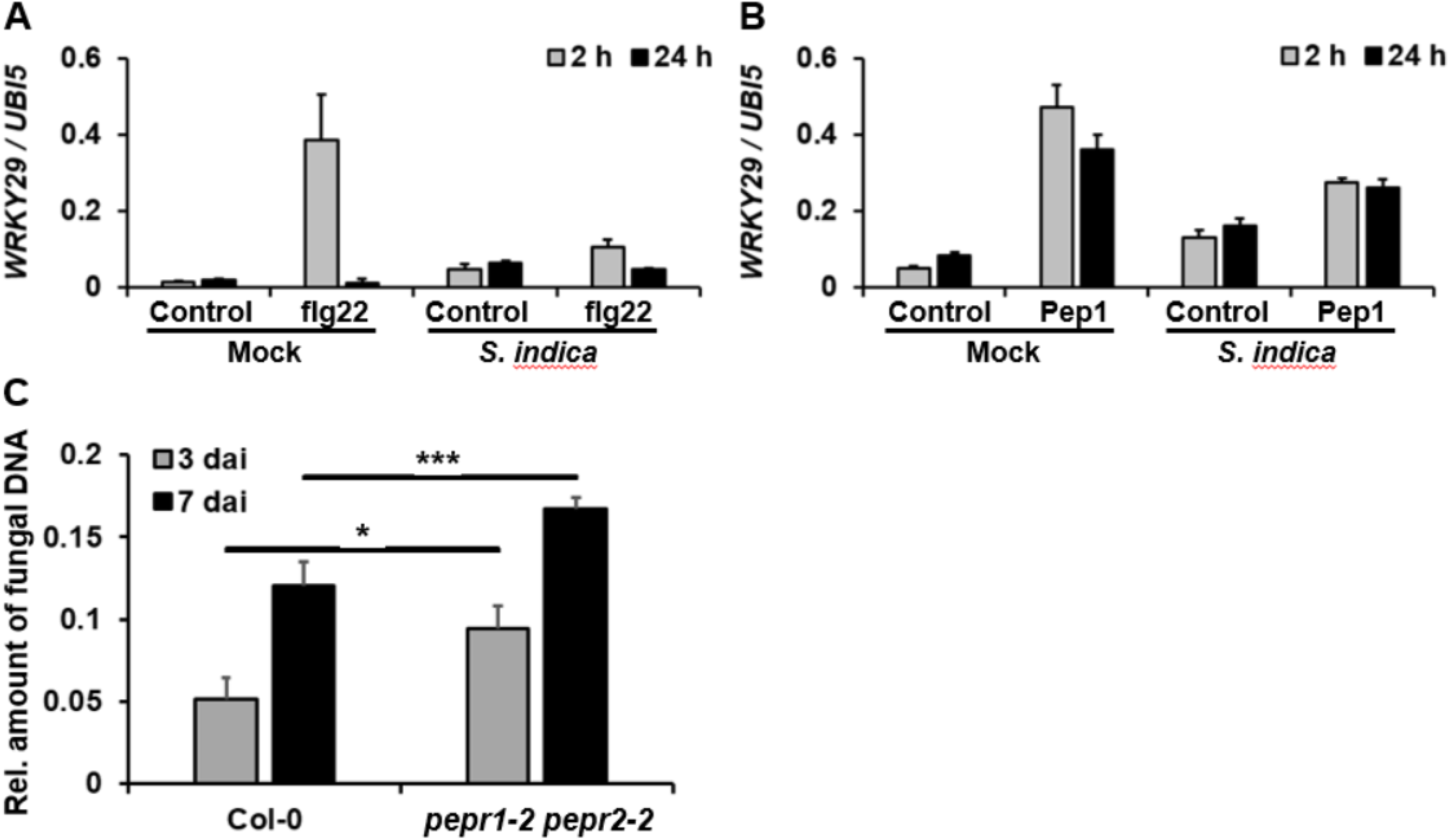
The ability of *Serendi ita indica* to suppress flg22 but not Pep1-triggered immune responses in roots. **(A, B)** S. *indica* inhibits flg22 but not Pep1 induction of PTI marker gene *WRKY29.* h, hours after flg22 or Pep1 treatment. **(C)** Improved root colonisation of *pepr1 pepr2* mutants by S. *indica.* dai, days after inoculation.

**Supplemental Figure 2.**
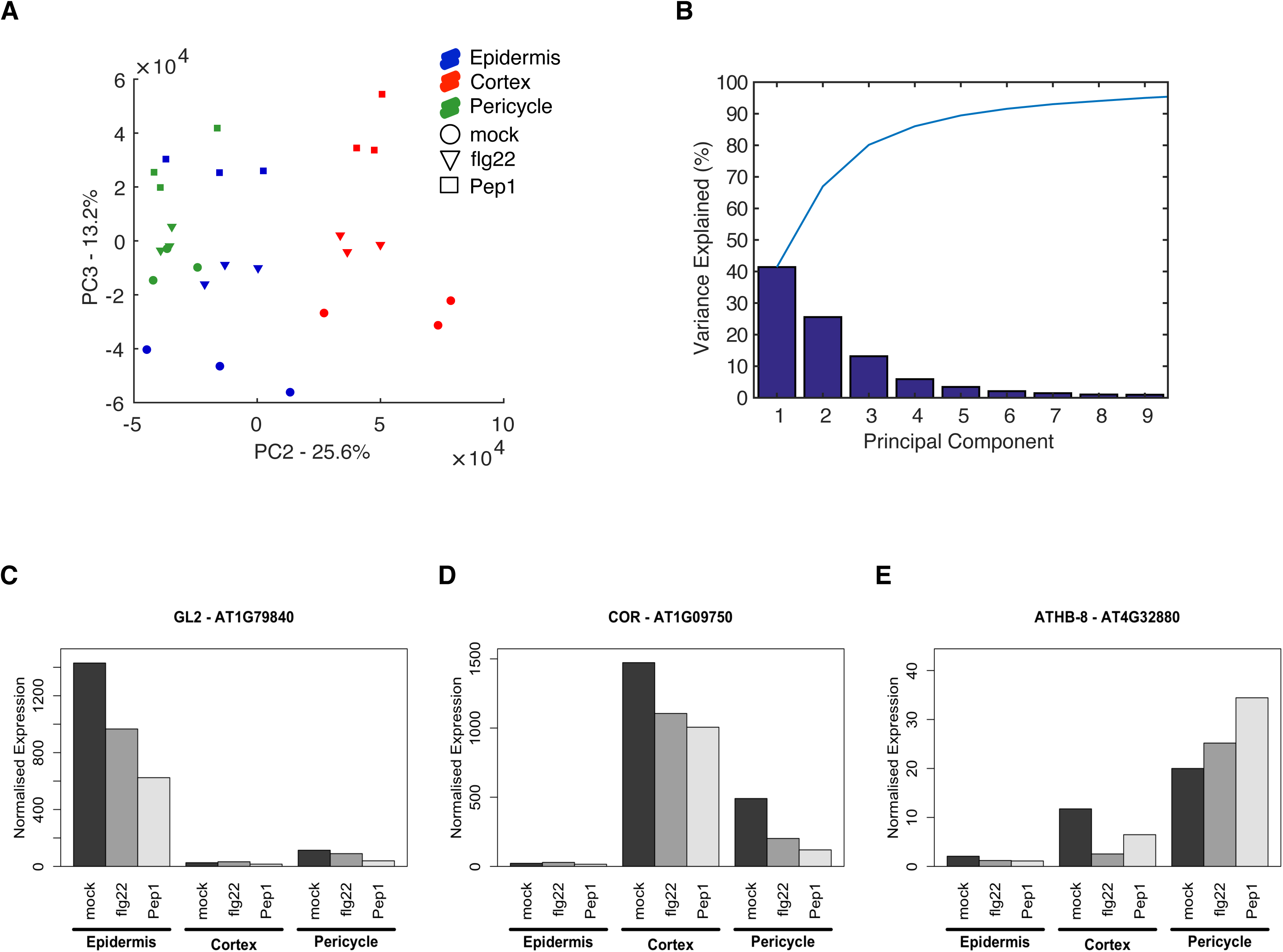
Assessment of RNA-seq data quality. **(A)** Plotting PC1 vs PC3 demonstrates the separation of samples based on cell type and immune elicitors. **(B)** Scree plot showing the percentage of variance explained by the top 10 PCs. **(C) (D) and (E)** Normalised RNA-seq expression data for cell-type markers, (GL2, CORTEX and ATHB-8 respectively).

**Supplemental Figure 3.**
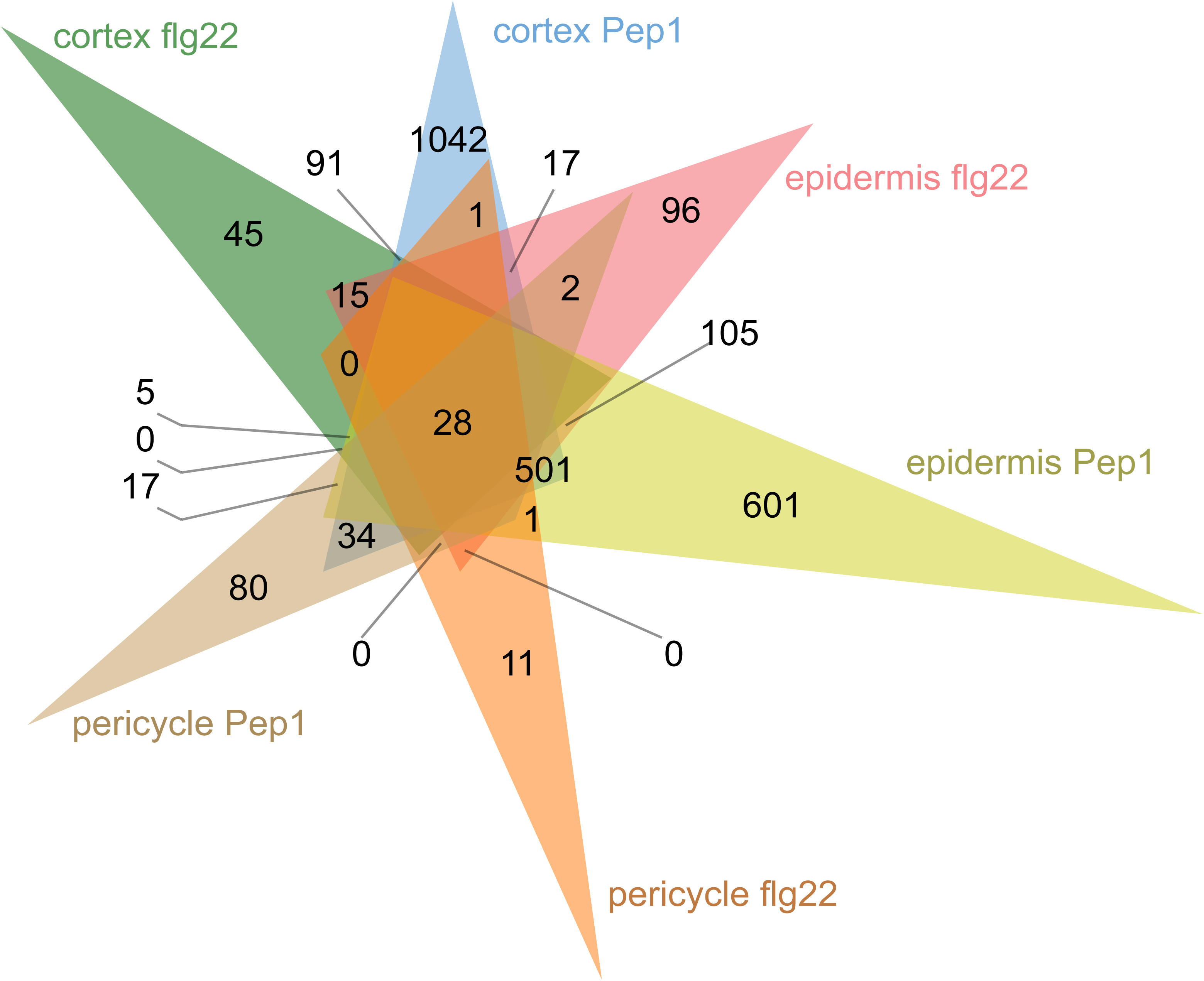
6-set Venn diagram detailing all overlaps between sets of DEGs for the two treatments in three cell-types revealing that 28 genes are differentially expressed in all three cell types following flg22 or Pep1 treatment.

**Supplemental Figure 4.**
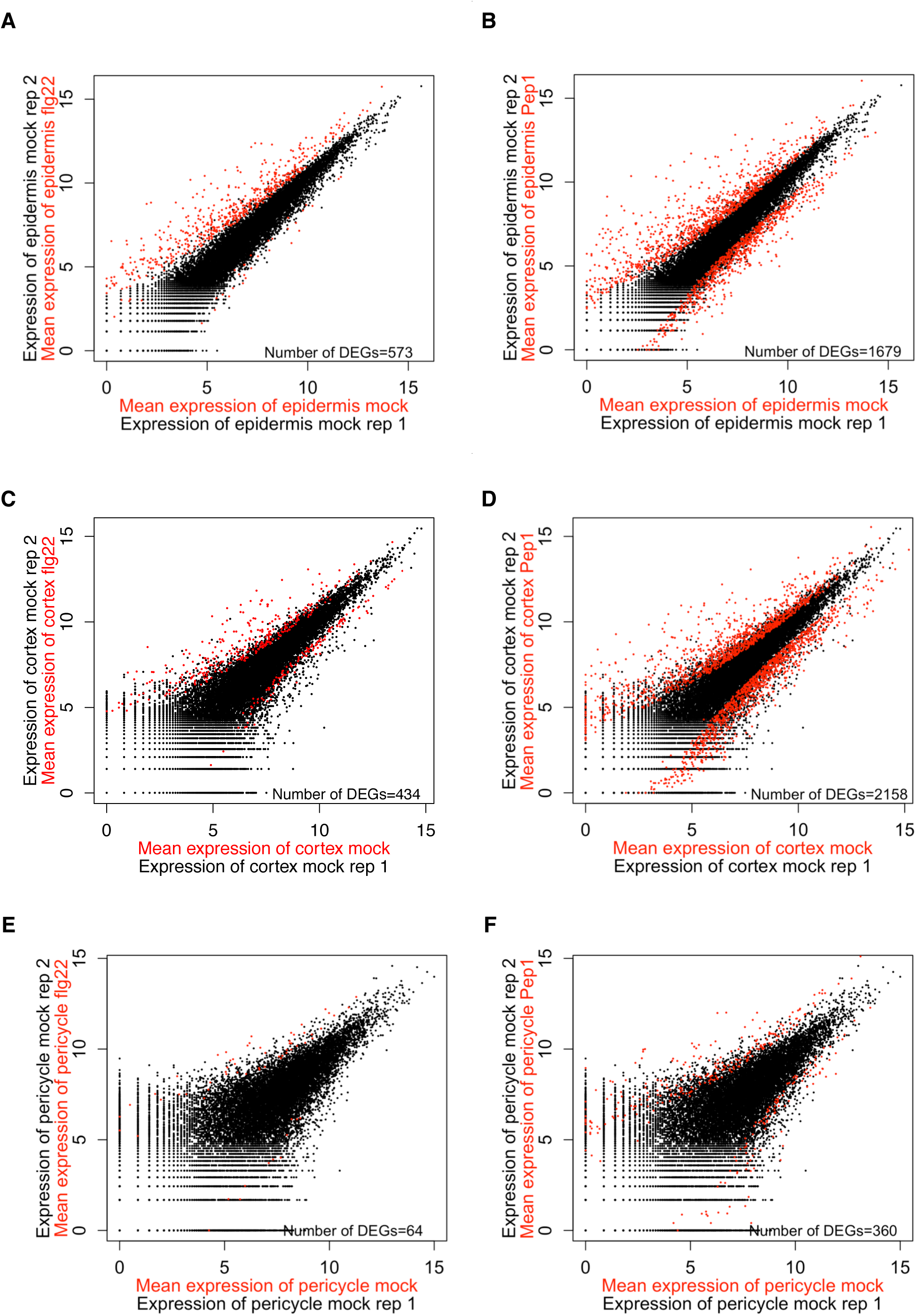
Visual inspection of differential gene expression. **(A) and (B):** Average log2 read counts of DEGs (flg22-vs mock-treated, (A), Pep1-vs mock-treated (B) detected by DESeq2 (red) overlaid over replicate plots showing the expression of all genes in two representative mock–treated epidermis replicates (black). **(C) and (D):** As above for flg22- and Pep1-treated replicates in the cortex ((C) and (D), respectively). **(E) and (F):** As above for flg22- and Pep1-treated replicates in the pericycle ((E) and (F), respectively).

**Supplemental Figure 5.**
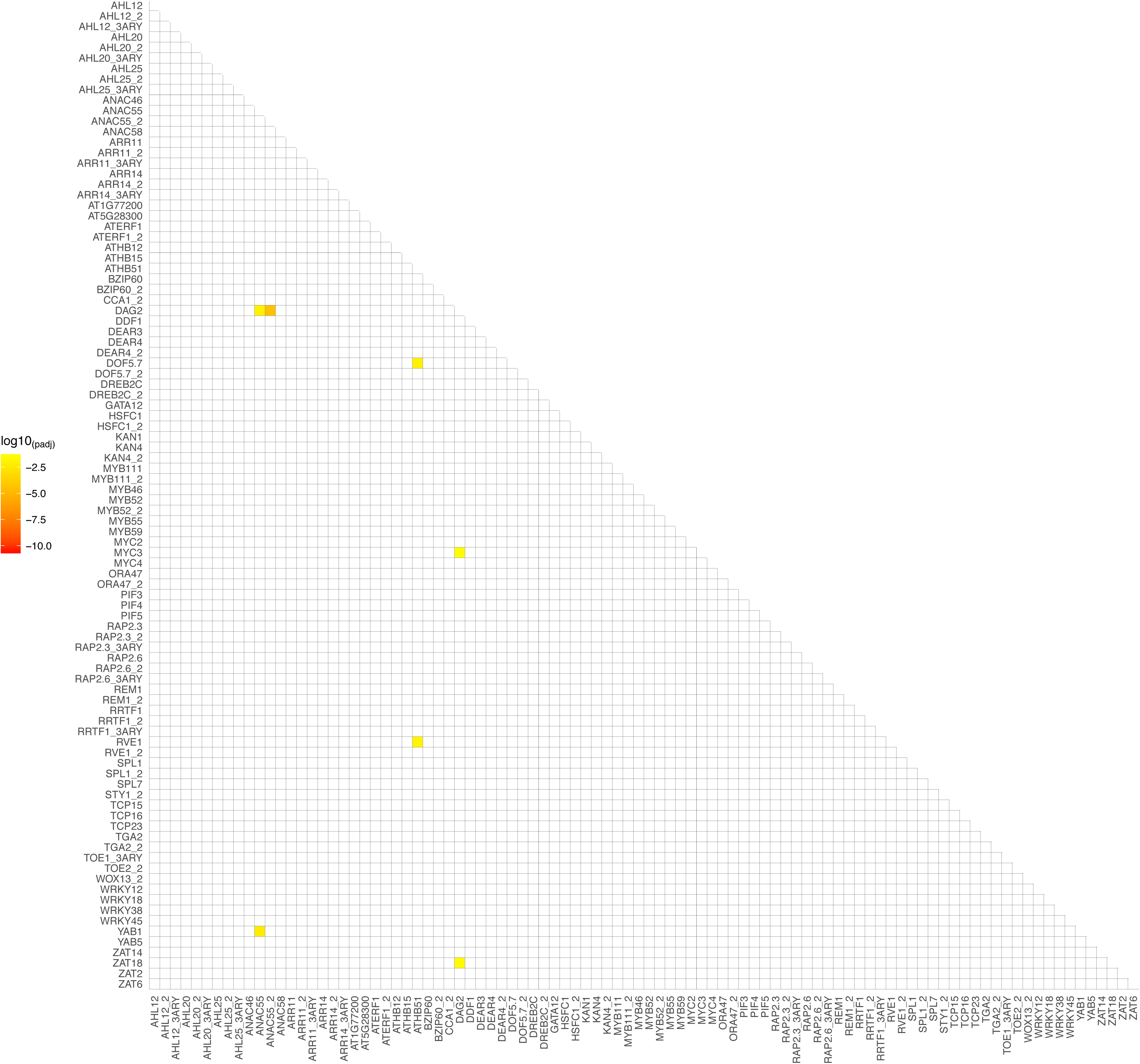
Heatmap representing p-values for enrichment of motif pairs in pericycle identity genes.

**Supplemental Figure 6.**
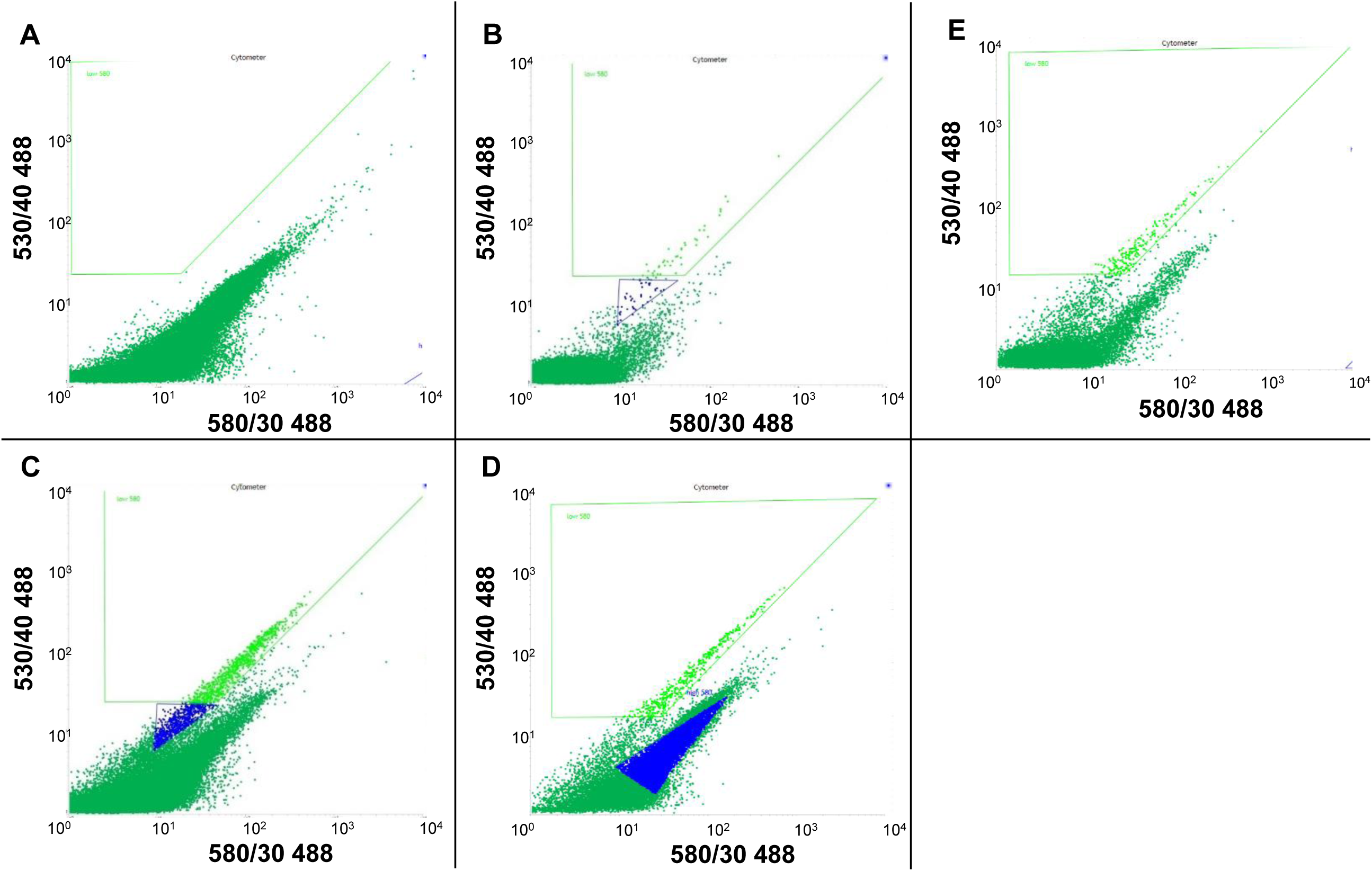
Representative FACS dot plots of the output from the 580/30 nm vs. the 530/40 nm bandpass filters. (A) Dot plot from Col-0 wildtype root protoplasts. (B-D) Dot plots from root-derived protoplasts from *pGL2:GFP* (B) *pCortex:GFP* and (C) *E3547* lines (D), respectively. During pre-experiments, different cell populations were collected to confirm presence/absence of green fluorescing cells microscopically (green and blue sorting gates). GFP-containing cells were located in the high 530 / low 580 populations. (E) Dot plot from *E3547* root protoplasts showing a sorting gate typically used in experiments to collect GFP-expressing protoplasts (green sorting gate).

## Supplemental Tables

**Supplemental Table 1.** Alignment statistics table based on output from htseq-count, explanation of columns can be found in htseq-count documentation at http://htseq.readthedocs.io/en/master/count.html

| Sample Name | Total Reads | mapped | No feature | ambiguous | Too low aQual | Not aligned | Alignment not unique | Fraction Unique Mapped |
| --- | --- | --- | --- | --- | --- | --- | --- | --- |
| WTCHG_125416_01 | 37936714 | 12787565 | 4093934 | 273829 | 0 | 0 | 32840287 | 33.70762 |
| WTCHG_125416_03 | 35029506 | 11652017 | 3788246 | 251581 | 0 | 0 | 29967974 | 33.26344 |
| WTCHG_125416_05 | 31334271 | 11378705 | 3059755 | 270834 | 0 | 0 | 24317704 | 36.31393 |
| WTCHG_129187_01 | 33036688 | 12753614 | 3151126 | 275370 | 0 | 0 | 28562074 | 38.6044 |
| WTCHG_129187_03 | 30640913 | 13289077 | 2555283 | 301281 | 0 | 0 | 23025575 | 43.37037 |
| WTCHG_129187_05 | 35134352 | 16675932 | 2645420 | 420388 | 0 | 0 | 22483846 | 47.46333 |
| WTCHG_129187_07 | 18770981 | 8309005 | 1750941 | 193046 | 0 | 0 | 14607485 | 44.26516 |
| WTCHG_129189_01 | 19821185 | 6775734 | 1968617 | 145809 | 0 | 0 | 16761463 | 34.1843 |
| WTCHG_129189_03 | 29569388 | 10700395 | 2799599 | 253587 | 0 | 0 | 24404250 | 36.18741 |
| WTCHG_129189_05 | 40014033 | 17564147 | 3338350 | 444268 | 0 | 0 | 27572569 | 43.89497 |
| WTCHG_129189_07 | 18831093 | 6654842 | 2154230 | 152612 | 0 | 0 | 16384479 | 35.33965 |
| WTCHG_129190_01 | 17092276 | 6915144 | 1710679 | 163863 | 0 | 0 | 14135806 | 40.45771 |
| WTCHG_129190_03 | 29240453 | 12110720 | 2895815 | 272652 | 0 | 0 | 23307571 | 41.41769 |
| WTCHG_129190_05 | 14936828 | 4936480 | 1722336 | 106923 | 0 | 0 | 13088899 | 33.04905 |
| WTCHG_129190_07 | 49209619 | 16040782 | 5944501 | 346962 | 0 | 0 | 42657375 | 32.59684 |
| WTCHG_131167_01 | 39127642 | 15036208 | 3509683 | 312311 | 0 | 0 | 33383597 | 38.42861 |
| WTCHG_131167_03 | 38637972 | 15850020 | 3259279 | 340108 | 0 | 0 | 31342063 | 41.02187 |
| WTCHG_131167_05 | 22346606 | 9665412 | 1871242 | 218768 | 0 | 0 | 16878898 | 43.25226 |
| WTCHG_203594_01 | 21806985 | 13680959 | 1351854 | 313391 | 0 | 0 | 8448254 | 62.73659 |
| WTCHG_203594_03 | 16606295 | 10233049 | 1106020 | 259778 | 0 | 0 | 6271995 | 61.62151 |
| WTCHG_203594_05 | 15128307 | 8843596 | 1100941 | 218599 | 0 | 0 | 6573581 | 58.45727 |
| WTCHG_203594_07 | 15672958 | 8711915 | 1188294 | 190475 | 0 | 0 | 7431993 | 55.58565 |
| WTCHG_203594_10 | 24230010 | 16017526 | 1335194 | 372547 | 0 | 0 | 7575272 | 66.10615 |
| WTCHG_203839_01 | 33380472 | 10609615 | 1035691 | 243146 | 0 | 0 | 6153122 | 31.7839 |
| WTCHG_203839_04 | 55203984 | 18302544 | 1561382 | 430497 | 0 | 0 | 8635572 | 33.15439 |
| WTCHG_203839_06 | 36293432 | 7479295 | 1814053 | 136844 | 0 | 0 | 13597890 | 20.60785 |
| WTCHG_203839_08 | 39814096 | 12477517 | 1286017 | 275089 | 0 | 0 | 7869031 | 31.33945 |

**Supplemental Table 2.**
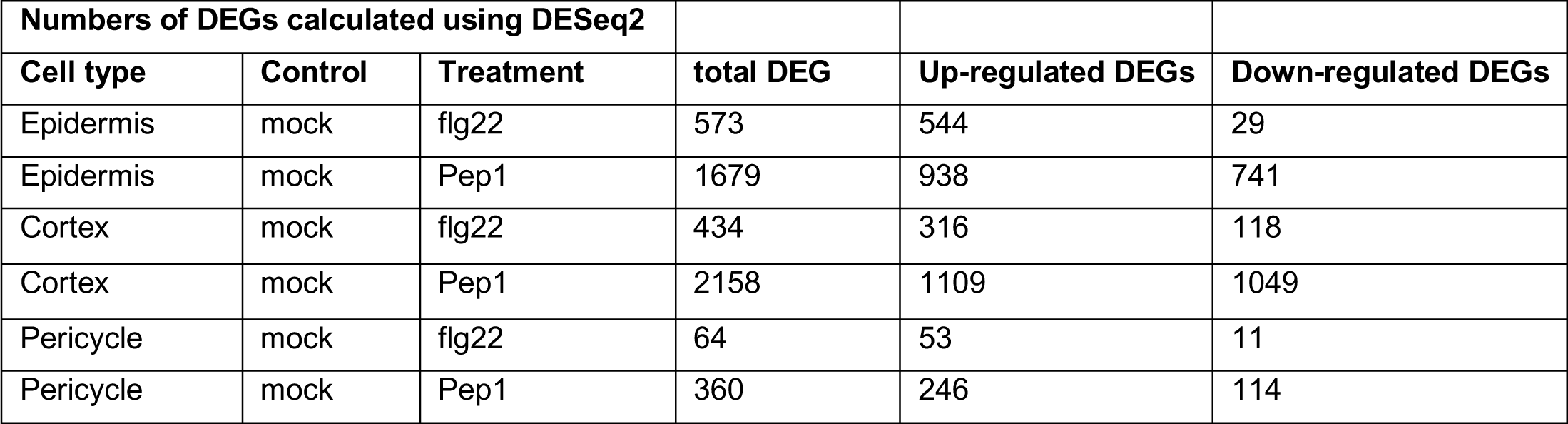
Numbers of DEGs responding to flg22 and Pep1 in three root cell types.

**Supplemental Table 3.** Genes differentially expressed in all three cell types in response to both flg22 and Pep1.

| ATG | names | description |
| --- | --- | --- |
| AT1G14540 | PER4 | Peroxidase superfamily protein |
| AT2G43620 | NA | Chitinase family protein |
| AT1G14550 | NA | Peroxidase superfamily protein |
| AT5G39580 | NA | Peroxidase superfamily protein |
| AT1G55450 | NA | S-adenosyl-L-methionine-dependent methyltransferases superfamily protein |
| AT1G21120 | IGMT2 | O-methyltransferase family protein |
| AT2G47550 | NA | Plant invertase/pectin methylesterase inhibitor superfamily |
| AT1G02930 | ATGST1 /<br>ATGSTF3 | glutathione S-transferase 6 |
| AT2G38870 | NA | Serine protease inhibitor; potato inhibitor I-type family protein |
| AT4G08850 | NA | Leucine-rich repeat receptor-like protein kinase family protein |
| AT4G01700 | NA | Chitinase family protein |
| AT1G02920 | ATGST11 /<br>ATGSTF7 | glutathione S-transferase 7 |
| AT1G02360 | NA | Chitinase family protein |
| AT5G48540 | NA | receptor-like protein kinase-related family protein |
| AT1G21110 | IGMT3 | O-methyltransferase family protein |
| AT1G21130 | IGMT4 | O-methyltransferase family protein |
| AT1G69920 | ATGSTU12 /<br>GSTU12 | glutathione S-transferase TAU 12 |
| AT3G53180 | NodGS | glutamate-ammonia ligases; catalytics; glutamate-ammonia ligases |
| AT3G10720 | NA | Plant invertase/pectin methylesterase inhibitor superfamily |
| AT1G52200 | NA | PLAC8 family protein |
| AT2G20960 | pEARLI4 | *Arabidopsis* phospholipase-like protein (PEARLI 4) family |
| AT1G07160 | NA | Protein phosphatase 2C family protein |
| AT3G51920 | ATCML9 / CAM9 | calmodulin 9 |
| AT5G64120 | AtPRX71 /<br>PRX71 | Peroxidase superfamily protein |
| AT5G67340 | NA | ARM repeat superfamily protein |
| AT2G39400 | NA | alpha/beta-Hydrolases superfamily protein |
| AT2G40180 | ATHPP2C5 /<br>PP2C5 | phosphatase 2C5 |
| AT3G19010 | NA | 2-oxoglutarate (2OG) and Fe(II)-dependent oxygenase superfamily protein |

**Supplemental Table 4.**
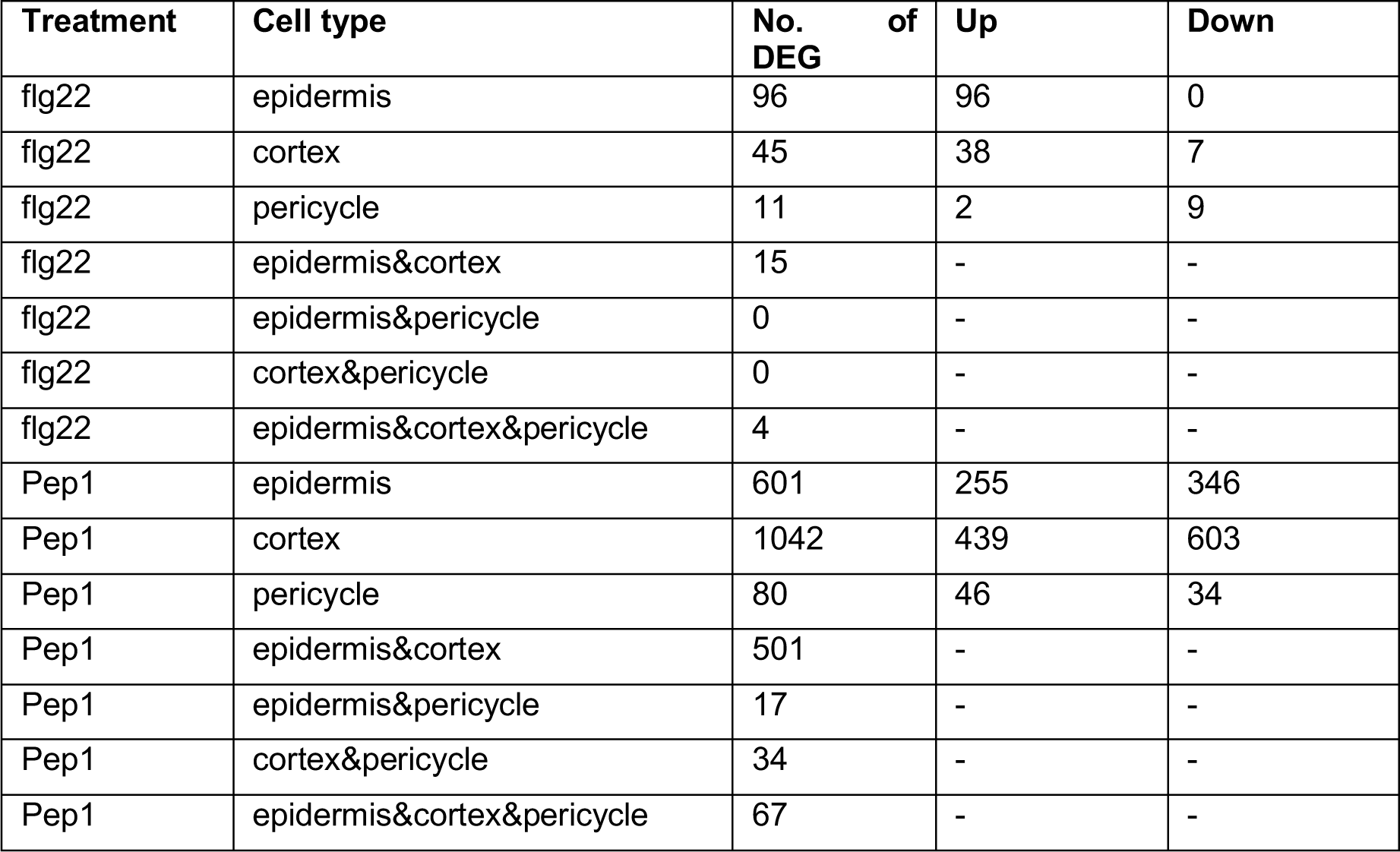
Numbers of DEGs responding specifically to either flg22 or Pep1 in one or more cell types.

**Supplemental Table 5.**
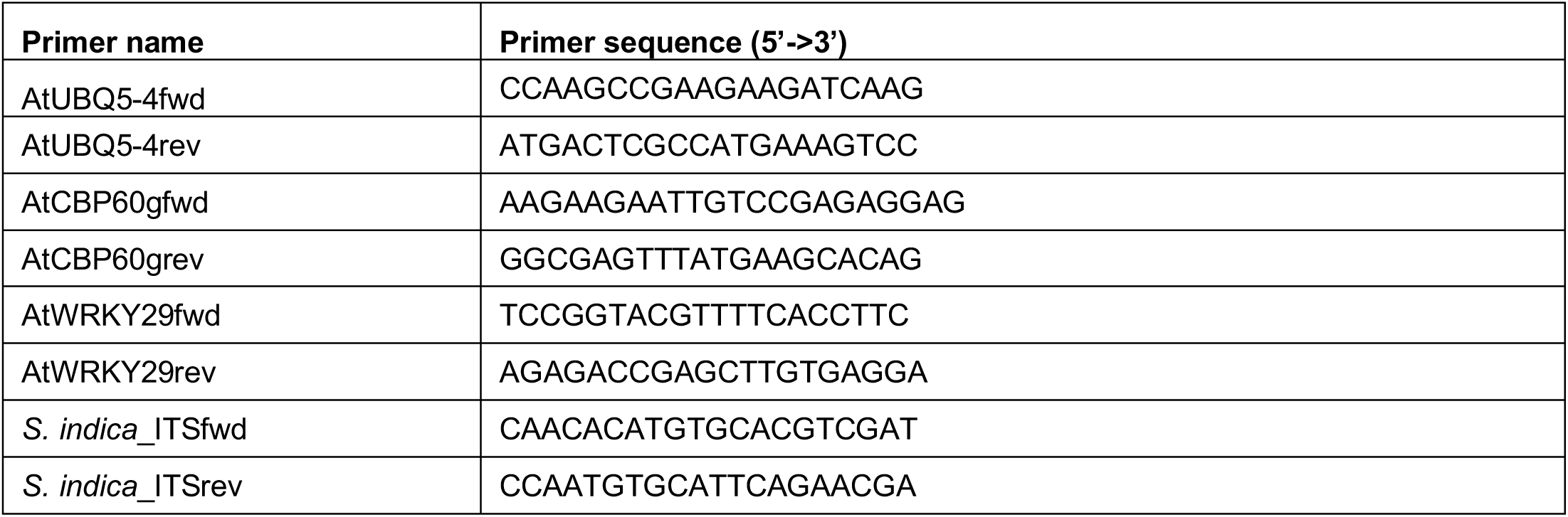
Primer sequences

